# Ex vivo ^1^H-MRS brain metabolic profiling in a two-hit model of schizophrenia-related alterations: effects of prenatal immune activation and peripubertal stress

**DOI:** 10.1101/641761

**Authors:** Roberto Capellán, Mario Moreno, Javier Orihuel, David Roura-Martínez, Marcos Ucha, Emilio Ambrosio, Alejandro Higuera-Matas

## Abstract

Prenatal infections are environmental risk factors for neurodevelopmental disorders. In addition, traumatic experiences during adolescence in individuals exposed to infections during gestation could increase the risk of schizophrenia. Given the tremendous burden that these diseases pose on societies when they fully manifest, it is of the most crucial importance to discover potential markers of the disease in its early stages or even before its onset, so that therapeutic strategies may be implemented. In the present study, we combined a well-characterized two-hit model of schizophrenia-related symptoms with proton magnetic resonance spectroscopy to discover potential biomarkers. To this end, we i.p. injected 100 μg/kg/ml of lipopolysaccharide (LPS) or saline on gestational days 15 and 16 to pregnant rats. Their male offspring were then subjected to five episodes of stress or handling on alternate days during postnatal days (PND) 28 to 38. Once the animals reached adulthood (PND70), we evaluated prepulse inhibition (PPI). At PND90, we performed an ex vivo proton magnetic resonance spectroscopy study in the cortex and striatum. While we did not detect alterations in PPI at the age tested, we found neurochemical disturbances induced by LPS, stress or (more interestingly) their interaction. LPS decreased glucose levels in the cortex and striatum and altered glutamate, glutamine and N-acetylaspartate levels. Glutamate and glutamine levels in the left (but not right) striatum were differentially affected by prenatal LPS exposure in a manner that depended on stress experiences. These results suggest that alterations in the glutamate cycle in the striatum may predate the full emergence of disorders of the schizophrenic spectrum and could be used as early markers of the disease.

## 1. INTRODUCTION

Several epidemiological studies have shown that maternal infection during pregnancy or postnatal exposure to traumatic experiences may precipitate the onset of certain diseases such as schizophrenia (Brown and Derkits, 2010; Fisher et al., 2014; Kentner et al., 2018; Rössler et al., 2014), autism (Lee et al., 2015; Zerbo et al., 2015) or bipolar disorder (Canetta et al., 2014; Parboosing et al., 2015). In the case of schizophrenia, the risk for the disease is seven-fold when the infection occurs in the first trimester of gestation (Brown et al., 2004). This maternal immune activation (MIA) induces cytokine secretion in the pregnant mother. Some of these released cytokines (such as IL-6) are capable of crossing the placental barrier, as well as the fetal blood-brain barrier, causing a marked impact on offspring neurodevelopment (Boksa, 2010). MIA can be induced and replicated in animal models by several agents; however, the most commonly used are polyinosinic-polycytidylic acid (Poly I:C) and lipopolysaccharide (LPS) (Boksa, 2010). The latter is a structural component of the outer membrane of Gram-negative bacteria and is recognized by Toll-like receptors 4 (TLR4) located in the plasma membrane of various cell types, such as macrophages, dendritic cells or mast cells (Chandler and Ernst, 2017; Vaure and Liu, 2014). Its administration triggers an inflammatory response characterized by an increase of numerous cytokines, such as IL-1β, IL-6, TNFα, CXCL1, CXCL2, CXCL10, CCL2 and CCL7, as well as an increase in circulating corticosterone (Golan et al., 2005; Ortega et al., 2011; Urakubo et al., 2001). The impact of the immunological challenge on adult offspring depends on injection timing, as well as on the dose employed (Kentner et al., 2018). Concurrently to the humans scenario, administration of the immunogen early in gestation is associated with more profound deficits than those observed when exposure occurs at a more advanced gestational stage (Meehan et al., 2017).

Even though MIA models provide an invaluable tool to explore the developmental causes of (neuro)psychopathologies, it has been argued that they may be insufficient to fully capture the complex interplay between antenatal and postnatal aetiologies of developmental disorders. Recently, two and even three hit models have been developed to include this interaction of causes. Giovanoli and colleagues used a MIA model combined with a subchronic stress exposure during adolescence in mice showing that MIA itself was not able to unmask the whole set of symptoms that were detected when MIA and stress at puberty were combined (Giovanoli et al., 2013). By using a similar approach, Burt et al. found decreased NMDA receptor function in the hippocampus of rats exposed to LPS during gestation, a treatment that also rendered this structure more susceptible to the effects of stress (Burt et al., 2013).

Prepulse inhibition (PPI) impairments are typically found in schizophrenic patients (Braff et al., 1978, 2001; Kumari et al., 1999) but also in other pathologies such as Huntington’s Disease (Swerdlow et al., 1995), obsessive compulsive disorder (Ahmari et al., 2012; Hoenig et al., 2005; Swerdlow et al., 1993), Asperger’s syndrome (McAlonan et al. 2002), Klinefelter syndrome (van Rijn et al., 2011), Fragile-X syndrome (Frankland et al., 2004) and Tourette syndrome (Castellanos et al., 1996). PPI reductions have also been reproduced in animal models of symptoms of schizophrenia (Swerdlow, Braff, and Geyer 2000), which makes it a helpful tool when assessing therapeutic interventions or potential causal factors. As regards this, Deslauriers and co-workers reported impaired PPI in a two-hit animal model. Interestingly, they only found PPI impairments when prenatal immune activation was followed by adolescent stress exposure, although it must be noted that PPI was measured 24 h after the last stress exposure, that is, during the pubertal period (Deslauriers et al., 2013).

Proton magnetic resonance spectroscopy (^1^H-MRS) is also a useful tool for the study of these disorders (Stagg and Rothman, 2014; Wijtenburg et al., 2015). The technique is especially useful when identifying and quantifying concentration of a large number of metabolites in various brain areas, facilitating a more comprehensive understanding of the neurochemical disturbances that underlie central nervous system pathologies (Stagg and Rothman, 2014). The most consistent finding from ^1^H-MRS studies is the reduction in N-acetylaspartate levels that has been observed in the thalamus, as well as in the frontal and temporal lobes of schizophrenic patients (Brugger et al., 2011; Wijtenburg et al., 2015). Also, in the prodromal stage, glutamate levels are elevated in the striatum and decreased in the thalamus of at-risk individuals (Fuente-Sandoval et al., 2011; Stone et al., 2009). Cortical glutamate reductions have also been observed in schizophrenia patients as compared to healthy controls. In a very elegant study, Thakkar and coworkers found differences between a compound group composed of schizophrenia patients and first-degree relatives as compared to healthy controls, suggesting that this decreased cortical glutamate might be a marker of disease liability (Thakkar et al., 2017). Even though in vivo proton magnetic resonance provides invaluable information of the living brain, the constraints imposed by the in vivo situation such as the relatively low magnetic fields that can be applied make it troublesome to accurately measure specific metabolites that could be important markers of the disease. Higher magnetic fields can be employed in the ex vivo approach, allowing for the reliable identification of a wider array of potential biomarkers. Consequently, the aim of this study was to apply ex vivo 11.7T ^1^H-MRS in a two-hit animal model of schizophrenia-related alterations to a perform a comprehensive search of possible markers of risk for the disease in the cortex and striatum, two brain regions known to be affected in schizophrenic patients.

## 2. MATERIALS AND METHODS

### 2.1. ANIMALS

Experiments were performed on the male offspring of 14-week-old male and 12-week-old female Sprague-Dawley rats obtained from Charles River (France). Animals were kept in a temperature and humidity-controlled environment (23 °C/50–60%), artificial light (12 h/12 h light/dark cycle, lights on at 8 am), ad libitum access to food (commercial diet for rodents A04: Panlab, Barcelona, Spain) and tap water. Rats were housed in transparent Plexiglas cages (48.3 cm length x 26.7 cm width x 20.3 cm height). All the procedures performed were compliant with European Union guidelines for the care of laboratory animals (EU Directive 2010/63/EU governing animal experimentation) and approved by the Bioethics Committee of UNED.

### 2.2. DRUGS

We used lipopolysaccharides from Escherichia coli 0111: B4 (Sigma-Aldrich) dissolved in a saline sterile solution (0.9% sodium chloride) as immunogen.

### 2.3. EXPERIMENTAL DESIGN

Male and female rats were mated one week after arrival to the animal facility. Vaginal smears were taken daily from the breeder females, and pregnancy was determined by the presence of sperm in the vaginal smear (day 0 of pregnancy). LPS was intraperitoneally injected to pregnant rats at a dose of 100 μg/kg/ml on gestational days (GD) 15 and 16 (Boksa, 2010; Fortier et al., 2007; Wischhof et al., 2015). The control group consisted of pregnant rats submitted to the same treatment schedule with saline injection instead of LPS. We imposed a limit of 12 pups per dam, culling the animal surplus. We marked the pups according to their prenatal treatment with a tattoo in the paw and ensured that each damn had an equal number of LPS and saline-exposed pups. In doing so, we homogenized any potential effect of prenatal treatment on maternal behaviour that could affect the development of the offspring. Litters were left undisturbed until postnatal day (PND) 21 when they were weaned and grouped in sets of 2-3. Each set belonged to the same litter and treatment. Between PNDs 28 and 38, we exposed the male offspring to unpredictable stress (peripubertal stress. PUS). This stage of development is a critical period known to be highly sensitive to the disrupting effects of traumatizing events relevant to neuropsychiatric disorders (Giovanoli et al., 2013). The stress protocol included five distinct stressors, applied on alternate days: 1) Stress by agitation (30 minutes in an orbital shaker at 100rpm). 2) Stress by immobilization (45 minutes in a cylindrical restainer under bright light). 3) Water deprivation for 16 hours. 4) 10 minutes of a forced swimming session in a water tank of 40 cm high x 18 cm in diameter at a temperature of 22 ± 1°C and a depth of 30 cm. 5) Constant changes of the home cage (five cage changes, with new sawdust, during the dark cycle at random intervals). Non-stressed controls received handling by the same researcher and on the same days as the stressed subjects. This protocol was adapted from the work of Giovanoli et al. 2013.

### 2.4. BEHAVIORAL ASSESSMENT

#### 2.4.1 Prepulse inhibition of the acoustic startle response (PPI)

At PND70-73, PPI of the acoustic startle was measured in a non-restrictive Plexiglas cage (28 × 15 × 17 cm) containing a vibration-sensitive platform (Cibertec). Rats were habituated for 7 minutes with a background noise of 65 decibels (dB), which continued throughout the session. Animals were exposed to 6 pulse-alone trials at the beginning and the end of the session, to stabilize the startle response and to calculate habituation percentage (these pulses were not included in the PPI calculations). The session was composed of 35 different trials: ten 120 dB pulse-alone trials, five null trials with no stimulus and twenty pulses preceded by a prepulse of 69- or 77-dB intensity (4 or 12 dB above the background noise, respectively) with an interval of 30 or 120 ms (Santos-toscano et al., 2016). The duration of the test was 20 min approximately. Prepulse inhibition is expressed as the % PPI and calculated using the following formula: 1 – [startle amplitude on prepulse + pulse trial / mean startle amplitude on pulse-alone trials]) × 100. The percentage of habituation is expressed as 100 × [(Mean of first pulse-alone block – Mean of last pulse-alone block) /Mean of first 6 pulse alone block].

### 2.5. NEUROCHEMICAL DETERMINATIONS

#### 2.5.1 Ex vivo proton magnetic resonance spectroscopy (^1^H-MRS)

At PND92, an ex vivo ^1^H-MRS study was performed in the Biomedical Research Institute “Alberto Sols” (CSIC-UAM, Madrid, Spain). Before this, at PND90, the animals underwent an in vivo MRI to study changes in brain volume, connectivity, and metabolism that will be reported elsewhere. The animals were sacrificed under deep isoflurane anesthesia and the brain tissue was rapidly fixed using a microwave fixation system. The whole cortex and striata were then dissected out and analyzed on a Bruker Avance 11.7 Tesla spectrometer (Bruker BioSpin, Karlsruhe, Germany) equipped with a 4 mm triple channel ^1^H/^13^C/^31^P High-Resolution Magic Angle Spinning (HR-MAS) resonance probe. Samples (15mg) were introduced into a 50-μl zirconia rotor (4mm OD) with 50 μl D_2_O and spun at 5000Hz at 4°C to prevent tissue degradation. Two types of monodimensional proton spectra were acquired using a water-suppressed spin echo Carr-Purcell Meiboom-Gill (CPMG) sequence with 36ms and 144ms echo time and 128 scans. Data were collected into 64k data point using a 10kHz (20 ppm) spectral width and water presaturation during 2 s relaxation delay. Total acquisition time was 16 minutes. Spectra were automatically analysed using LCModel software (Provencher, 2001), 6.2-OR version (Oakville, ON; Canada). Only the peak concentrations obtained with a standard deviation lower than 20% were accepted.

### 2.5. STATISTICAL ANALYSIS

Data were analysed with IBM SPSS Statistics 24 for Windows. All results are expressed as the mean ± standard error of the mean (SEM) in the graphs or mean ± standard deviation (SD) in the tables. Outliers were identified by SPSS using the interquartile range criterion and a value of p < 0.05 was considered to represent a statistically significant difference. Square root, Neperian logarithmic and inverse transformations were applied when appropriate to correct the skewness in the distribution of the data and the lack of homogeneity of variances. PPI and ^1^H-MRS results were analyzed by two-way ANOVA considering “Prenatal immune activation” (saline or LPS) and “PUS exposure” (no stress or stress) as between-subject factors. We analysed interactions by simple effects analysis. The non-parametric Kruskal-Wallis H test was used if ANOVA assumptions were not met. F-value, effect sizes (partial eta square, η^2^_p_) and degrees of freedom are also reported when appropriate.

## 3. RESULTS

### 3.1 Behavioral outcomes

#### 3.1.1 Prepulse inhibition of the acoustic startle response was altered neither in LPS-treated animals nor in those exposed to peripubertal unpredictable stress

No significant effects were observed by prenatal immune activation or PUS exposure in any of the experimental conditions used in the PPI test (Figure 1). Negative values (reflecting prepulse facilitation) were not included in the analysis. Results obtained in the statistical tests are detailed in Table 1.

**Figure 1:**
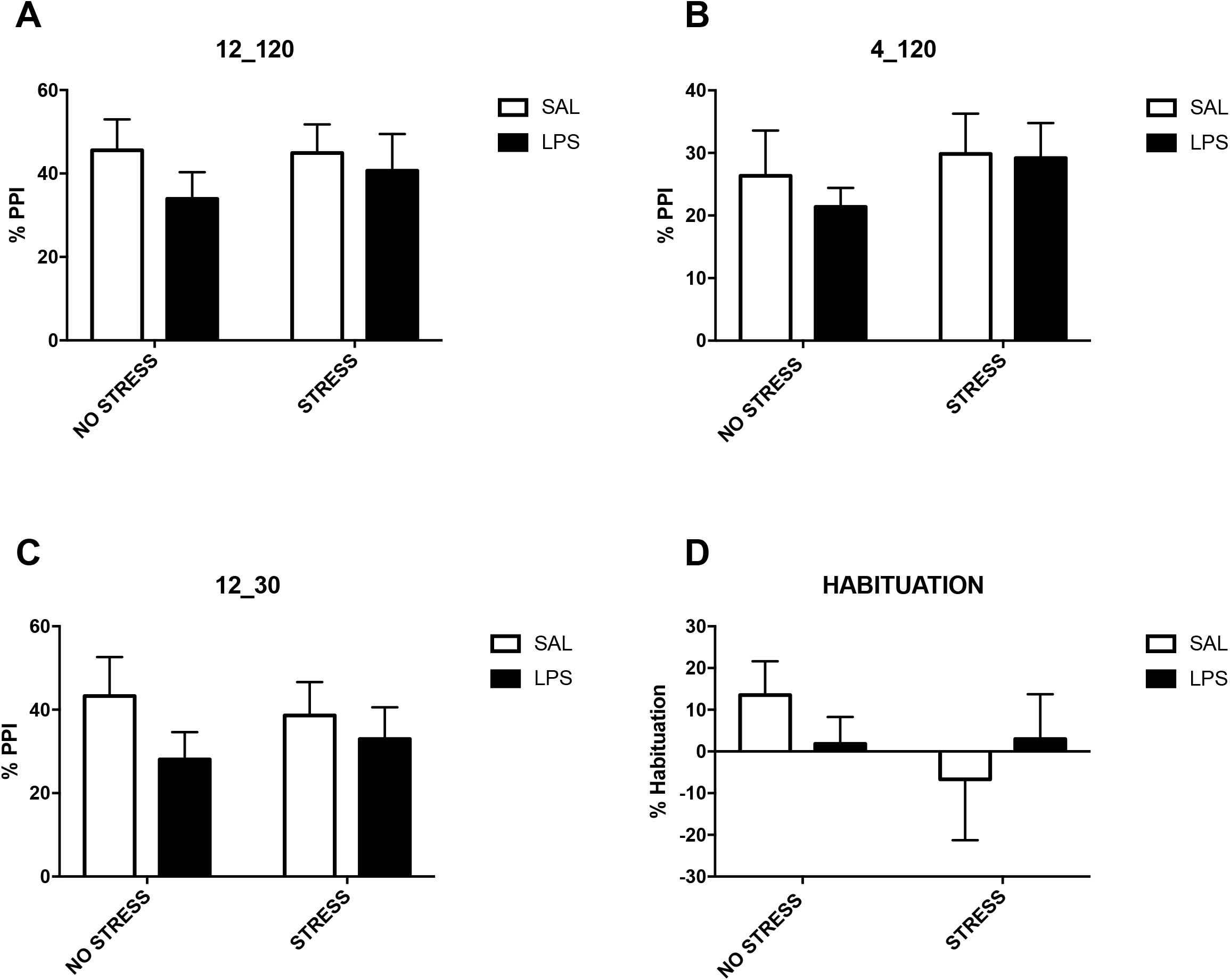
Effects of prenatal LPS treatment or PUS on % PPI and % Habituation. The figure shows % PPI at (A) 12 dB prepulse intensity and 120 ms interval (SAL+ NO STRESS: n=8; SAL + STRESS: n=8; LPS + NO STRESS: n=8; LPS + STRESS: n=8); (B) 4 dB prepulse intensity and 120 ms interval (SAL + NO STRESS S: n=8; SAL + STRESS: n=7; LPS + NO STRESS: n=7; LPS + STRESS: n=6); (C) 12 dB prepulse intensity and 30 ms interval (SAL + NO STRESS: n=8; SAL + STRESS: n=8; LPS + NO STRESS: n=7; LPS + STRESS: n=8). % Habituation was also calculated (D) (SAL + NO STRESS: n=8; SAL + STRESS: n=8; LPS + NO STRESS: n=8; LPS + STRESS: n=8).

**Table 1:** Summarized statistical results of the PPI test.

| PPI Condition | F-value and degrees of freedom | P-value |
| --- | --- | --- |
| 12_120 |  |  |
| Maternal immune activation | F (1, 28) = 1.15 | 0.292 |
| Peripubertal unpredictable stress | F (1, 28) = 0.167 | 0.585 |
| Interaction | F (1, 28) = 0.253 | 0.618 |
| 4_120 |  |  |
| Maternal immune activation | F (1, 24) = 0.217 | 0.723 |
| Peripubertal unpredictable stress | F (1, 24) = 0.889 | 0.355 |
| Interaction | F (1, 24) = 0.127 | 0.645 |
| 12_30 |  |  |
| Maternal immune activation | F (1, 27) = 1.665 | 0.207 |
| Peripubertal unpredictable stress | F (1, 27) = 0.000 | 0.988 |
| Interaction | F (1, 27) = 0.348 | 0.560 |
| Habituation to main stimulus |  |  |
| Maternal immune activation | F (1, 28) = 0.010 | 0,920 |
| Peripubertal unpredictable stress | F (1, 28) = 0.825 | 0,371 |
| Interaction | F (1, 28) = 1.04 | 0,316 |

### 3.2 ^1^H-MRS determinations

All metabolite concentrations were normalized to creatine + phosphocreatine levels [Cr+PCr], due to its relatively constant concentration in the brain. No statistically significant changes were detected in this peak across groups. Tables 1–8 of the supplementary material show the summarized statistical results of each metabolite and their mean ± SD in the left cortex, right cortex, left striatum and right striatum, respectively. The graphs show the most relevant results.

#### 3.2.1 Glucose levels were decreased bilaterally in the striatum and the left cortex of LPS-exposed animals

Prenatal immune activation diminished glucose levels in left (F_1,25_ = 8.586, p = 0.007, η^2^_p_ = 0.256) (Figure 2, A) and right (F_1,26_ = 12.402, p = 0.002, η^2^_p_ = 0.323) (Figure 2, B) striata, as compared to the saline groups. This effect was also observed in the left cortex (F_1,27_ = 5.855, p = 0.023, η^2^_p_ = 0.178) (Figure 2, C).

**Figure 2:**
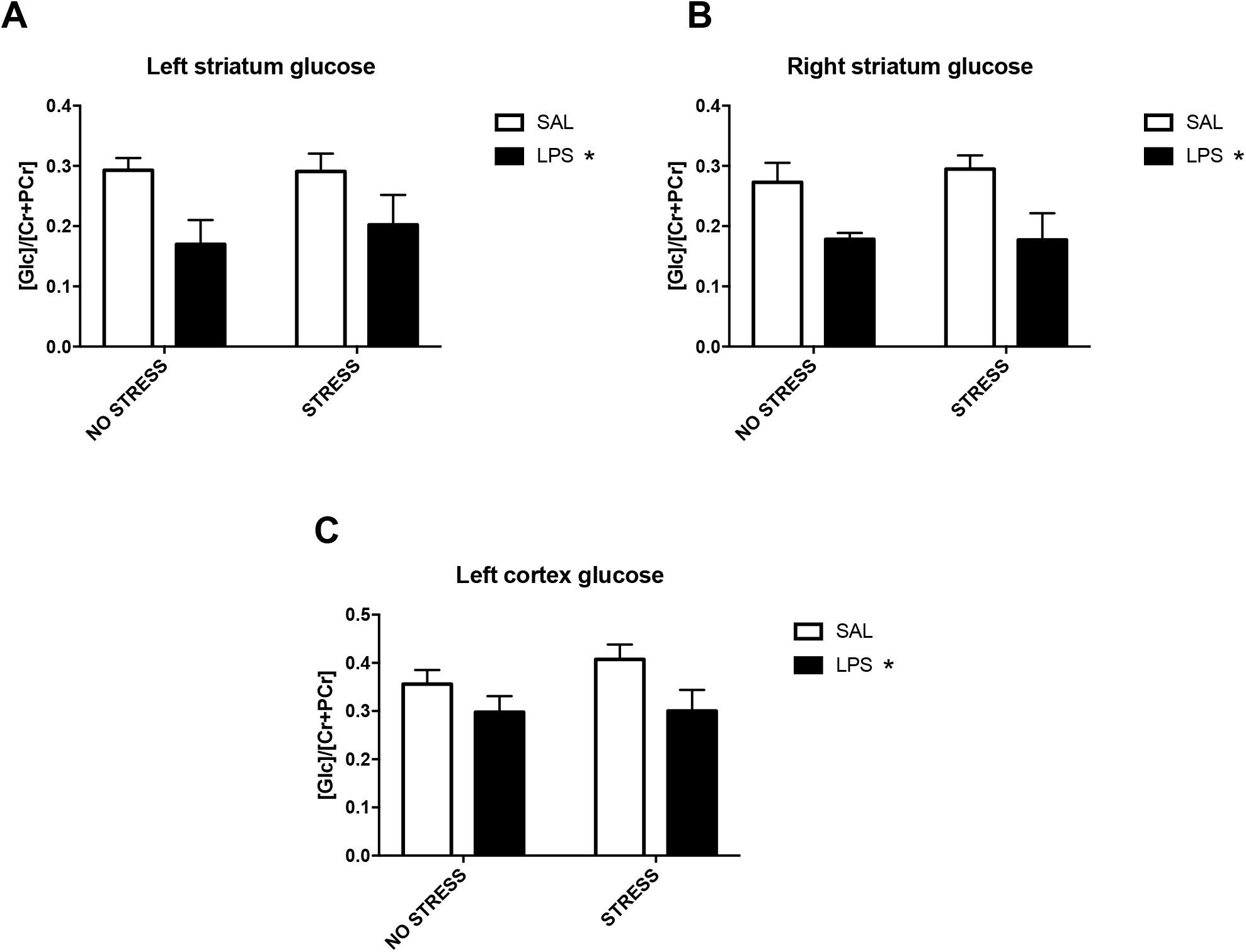
Normalised glucose levels in (A) left striatum (SAL + NO STRESS: n=7; SAL + STRESS: n=8; LPS + NO STRESS: n=7; LPS + STRESS: n=7), (B) right striatum (SAL + NO STRESS: n=8; SAL + STRESS: n=8; LPS + NO STRESS: n=7; LPS + STRESS: n=7) and (C) left cortex (SAL+ NO STRESS: n=8; SAL + STRESS: n=8; LPS + NO STRESS: n=8; LPS + STRESS: n=7). * p < 0.05 compared to the respective saline groups.

#### 3.2.2. Glutamate levels were increased in the left cortex and reduced in the left striatum of LPS-exposed animals. The combination with peripubertal stress increased glutamate levels in the striatum

Higher glutamate levels (F_1,27_ = 4.42, p = 0.045, η^2^_p_ = 0.141) were obtained in the left cerebral cortex of LPS-exposed animals (Figure 3, A). Moreover, a significant interaction was observed between prenatal immune activation and PUS exposure (F_1,27_ = 11.21, p = 0.002, η^2^_p_ = 0.293) in the left striatum. In this area, LPS-exposed animals showed lower glutamate levels in the absence of PUS (F_1,27_ = 5.454, p = 0.027). However, in its presence, glutamate levels were increased (F_1,27_ = 5.756, p = 0.024). Likewise, PUS increased glutamate levels only in LPS-exposed animals (F_1,27_ = 12.219, p = 0.002) (Figure 3, B).

**Figure 3:**
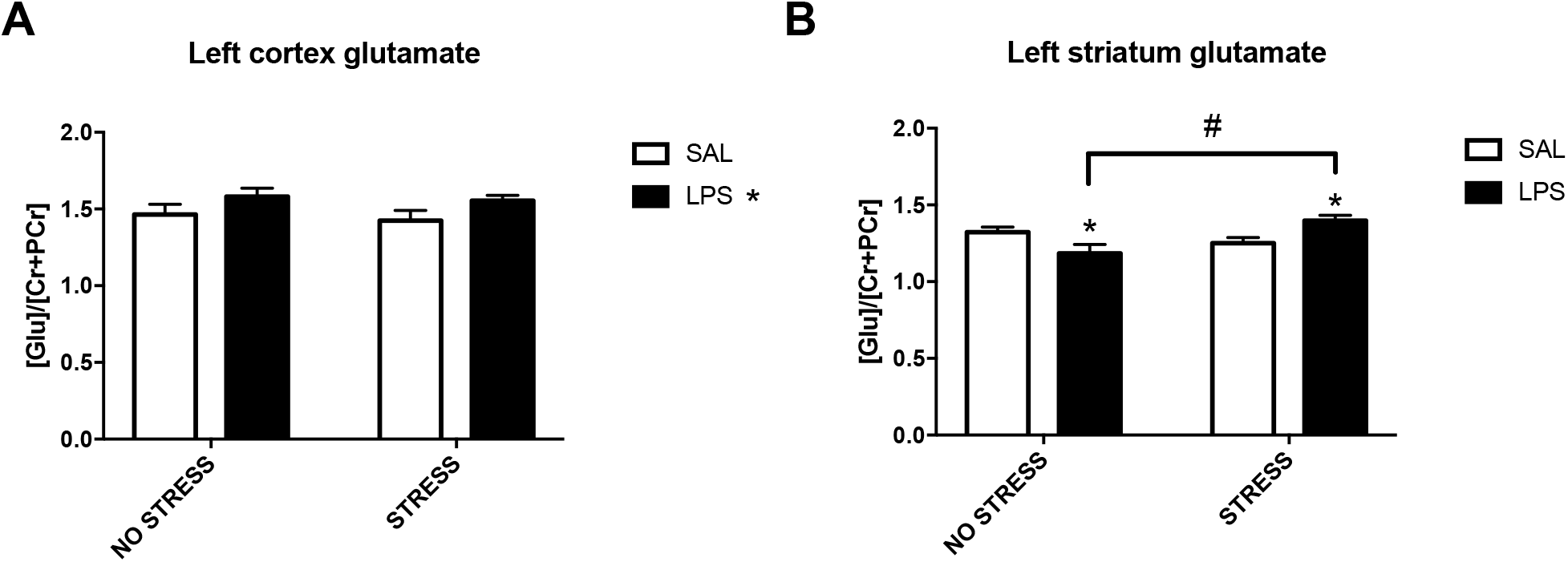
Normalised glutamate levels in left cortex (A) (SAL + NO STRESS: n=8; SAL + STRESS: n=8; LPS + NO STRESS: n=8; LPS + STRESS: n=7) and left striatum (SAL + NO STRESS: n=8; SAL + STRESS: n=8; LPS + NO STRESS: n=8; LPS + STRESS: n=7). * p < 0.05 compared to the respective saline groups. # Significant difference (p < 0.05) between groups.

#### 3.2.3. Glutamine levels were reduced in the left striatum by prenatal immune activation and PUS exposure

A significant interaction was found between prenatal immune activation * PUS exposure (F_1,27_ = 6.155, p = 0.02, η^2^_p_ = 0.186) in the levels of glutamine in the left striatum. In this area, LPS-exposed animals showed lower glutamine levels, in absence of PUS (F_1,27_ = 5.641, p = 0.025). This effect was also obtained in peripubertally-stressed animals, but only in the absence of prenatal immune activation (F_1,27_ = 4.411, p = 0.045) (Figure 4).

**Figure 4:**
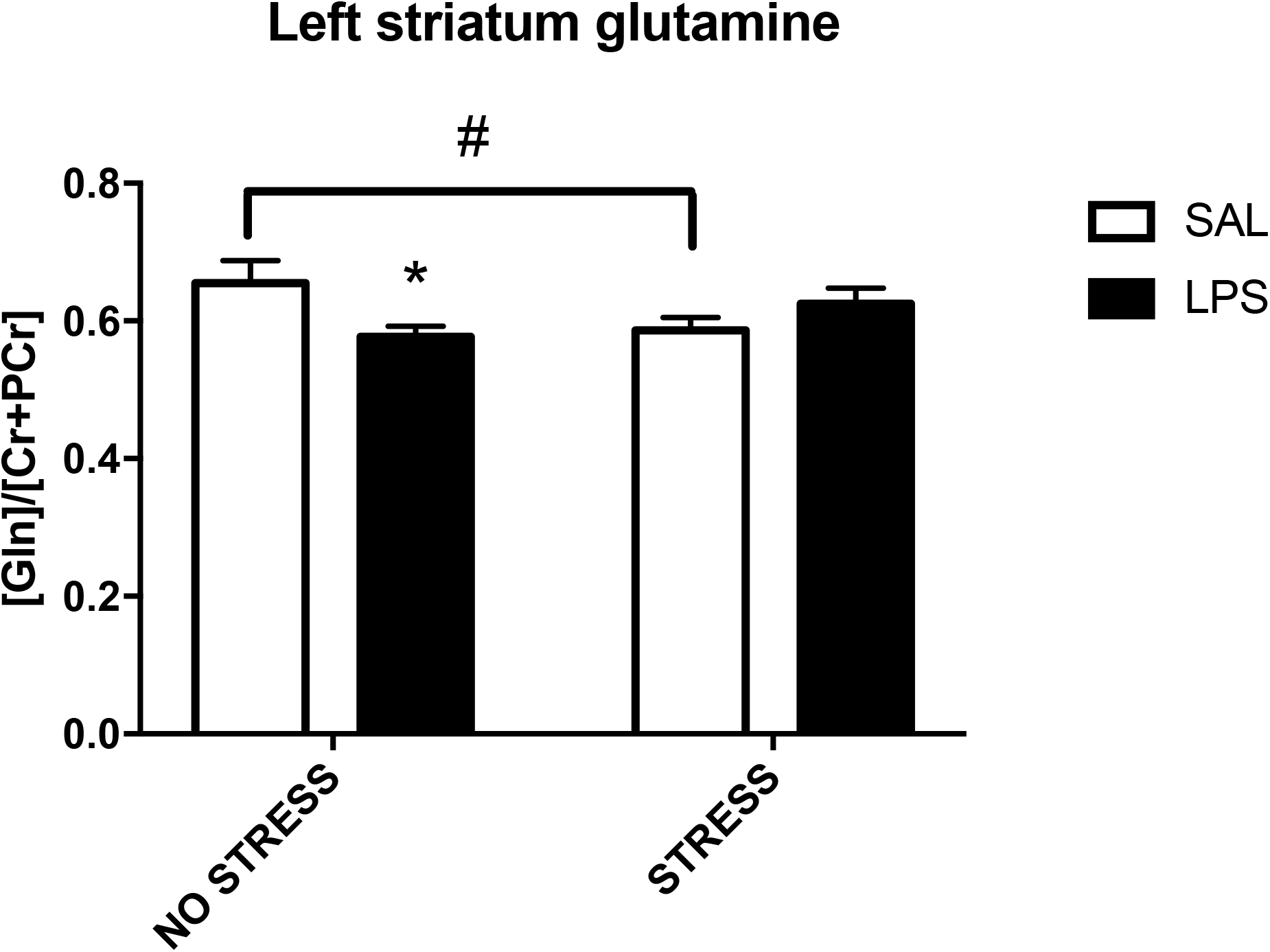
Normalised glutamine levels in left striatum (SAL + NO STRESS: n=8; SAL + STRESS: n=8; LPS + NO STRESS: n=8; LPS + STRESS: n=7). * p < 0.05 compared to the respective saline groups. # Significant difference (p < 0.05) between groups.

#### 3.2.4. *N*-acetylaspartate levels were reduced in the right cortex of LPS-exposed animals

LPS-exposed animals showed significantly lower N-acetylaspartate levels (F_1,27_ = 4.438, p = 0.045, η^2^_p_ = 0.141) in right cerebral cortex (Figure 5).

**Figure 5:**
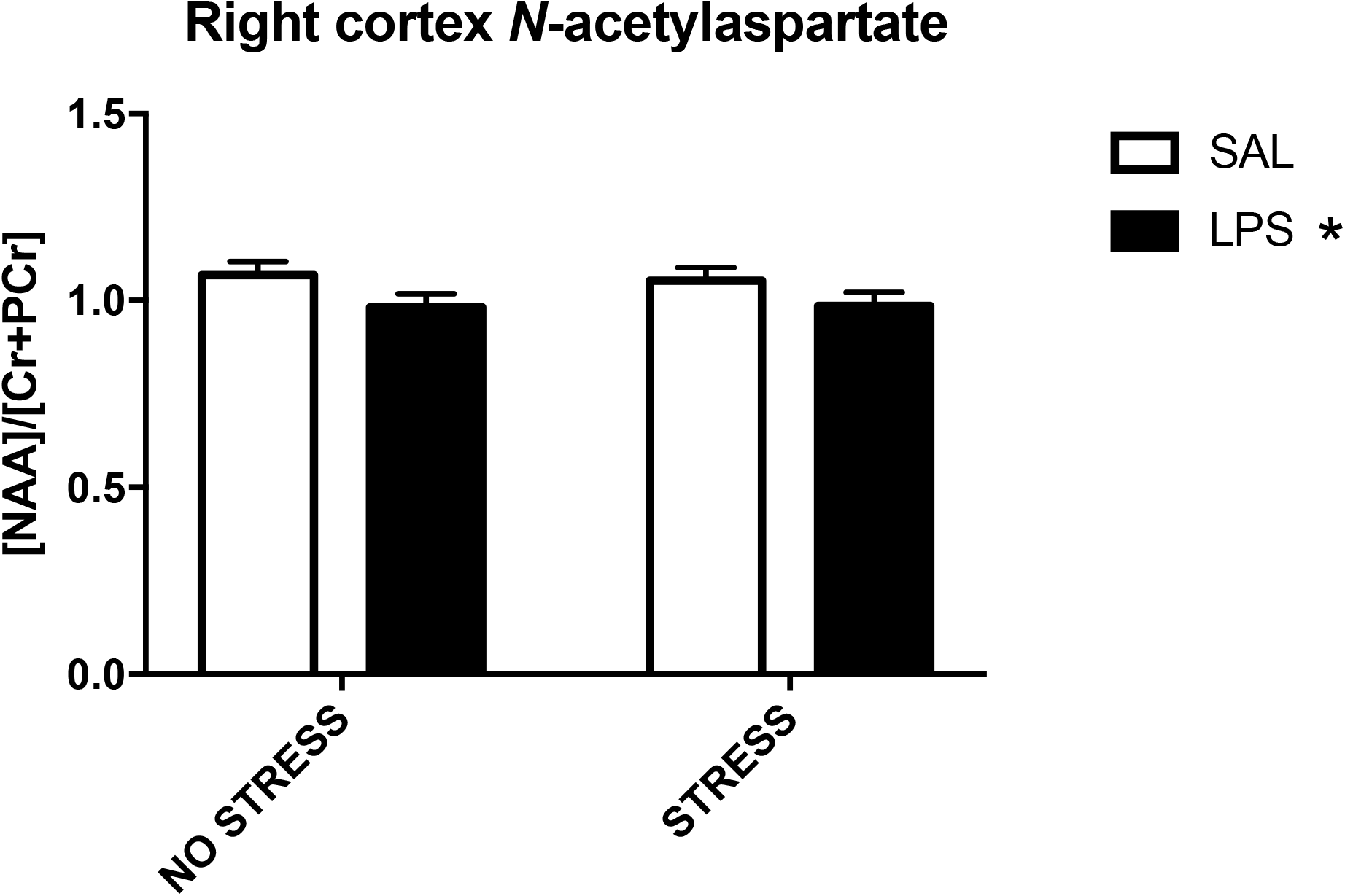
Normalised N-acetylaspartate levels in the right cerebral cortex (SAL + NO STRESS: n=8; SAL + STRESS: n=8; LPS + NO STRESS: n=8; LPS + STRESS: n=7). * p < 0.05 compared to the respective saline groups.

## 4. DISCUSSION

### 4.1 Prepulse inhibition of the acoustic startle response

Under our specific experimental conditions, we did not observe a significant effect of prenatal LPS administration, pubertal stress exposure, or their combination on sensorimotor gating in the PPI test. This absence of effects contrasts with the synergic effect between prenatal immune activation and pubertal stress reported by Giovanoli and coworkers (Giovanoli et al., 2013) who showed that PPI impairments in mice were only present when the animals were exposed to the two hits (prenatal immune activation with Poly I:C and pubertal stress) combined. Impaired PPI has consistently been documented in maternal immune activation models, especially those involving Poly I:C (see Meyer 2014 for a review) and also with LPS exposure (Fortier et al., 2007; Romero et al., 2010; Santos-Toscano et al., 2016; Simões et al., 2018; Swanepoel et al., 2018; Waterhouse et al., 2018; Wischhof et al., 2015) although some variables such as the species used (Imai et al., 2018), the prenatal period at which exposure occurs (Waterhouse et al., 2018) or the age of testing (Basta-Kaim et al., 2012) influence the magnitude or actual presence of PPI impairments. In these experiments, we chose to study PPI at a slightly earlier period of development as compared to a previous study by our group (PND 70-73 here, as compared to PND72-77 in Santos-Toscano et al.) and did not include females. Moreover, as observed in Figure 1, PUS seems to be counteracting LPS effects to some extent. Therefore, a stronger effect of LPS exposure might be needed to obtain a main effect of the maternal immune activation or an interaction between both factors. These results could also be interpreted as prodromal in the sense that PPI impairments would be incubating and only after acute stress or other developmental factors later in life, the deficits would be unmasked. In this context, the identification of early neural markers at this prodromal stage is especially relevant.

### 4.2 Neurochemical determinations

As a first step in the search for early markers of schizophrenia, it is essential to study as many candidate molecules as possible to identify potential biomarkers of the incipient disease. Ex vivo MRS is a suitable technique to attain this goal, but the results obtained should be validated in preclinical models with in vivo ^1^H-MRS and in human patients.

#### 4.2.1 Glucose

LPS-exposed animals showed lower glucose levels in the striatum, both in left and right hemispheres. In the cortex, this effect was only found in the left hemisphere. Given that glucose is the primary energetic substrate of the brain, we can speculate that a reduction in the glucose contents of these areas may be a consequence of a regional hypoactivation. In accordance with these results, PET studies in human schizophrenic patients have documented decreases in glucose metabolic rate in the striatum and the cortex (Hazlett et al., 2019) and, in animal models, rats with a subchronic exposure to MK-801 showed decreased glucose levels in the temporoparietal cortex, as assessed with ^13^C-MRS (Eyjolfsson et al., 2011). As regards this, glucose metabolism seems to be altered in the striatum of schizophrenic patients. Indeed, the levels of the β subunit of pyruvate dehydrogenase were lower and levels of pyruvate and lactate were significantly higher in brain samples containing the striatum from subjects with a diagnosis of schizophrenia, (Dean et al., 2016). As a whole, this altered corticostriatal function has been suggested to be part of the complex assembly of factors mediating liability for schizophrenia (Wagshal et al., 2014). Given that our glucose results were only associated to MIA and not the combination of MIA and PUS, we propose that these lower glucose levels are indeed suggestive of liability or vulnerability to the disorder rather than a marker of ultra-high-risk or disease onset.

#### 4.2.2 Glutamate, glutamine and N-acetylaspartate

We decided to group the discussion of these three metabolites in a single category due to their joint involvement in the glutamatergic system. Glutamate, in addition to its essential role in cellular metabolism, is the most abundant excitatory neurotransmitter. Glutamine has been suggested to be an informative marker of glutamatergic neurotransmission, as it is generated after synaptic glutamate is captured by astrocytes (Bak et al., 2006). N-acetylaspartate levels are related to neuronal density and functionality (Demougeot et al., 2001). Also, this molecule is a precursor for the biosynthesis of N-acetylaspartylglutamate, a neuropeptide with actions at NMDA and metabotropic glutamate receptors (Neale et al., 2000). We have found a fascinating pattern of results regarding glutamate and glutamine in the left striatum. While MIA per se was associated with a decrease in glutamate and glutamine levels, the combination of MIA with PUS significantly increased glutamate levels above those seen in stressed animals without MIA. As regards glutamine, the pattern was almost identical: while MIA alone resulted in decreased levels of this amino acid, the MIA + PUS tended to increase glutamine levels.

As stated before, while exposure to MIA decreased glutamate accumulation in the left striatum, exposure to both hits, a situation that is suggestive of either a high risk for the disease (more so if we consider the absence of overt signs of PPI impairment in our animals) or an initial onset, was associated with increased glutamate abundance in the region. This is in accordance to human studies where individuals at ultra-high-risk of schizophrenia and those with a first episode of psychosis that subsequently transitioned to the actual disease had higher levels of glutamate/glutamine in the (right) dorsal caudate nucleus (see Wijtenburg et al. 2015 for a review). Interestingly, this glutamatergic signature is no longer observable once the disease has reached chronicity (Tayoshi et al., 2009).

Glutamate levels were increased in the left cortex of rats exposed to the immune challenge during gestation. There is no consensus in the literature with regard to glutamate levels in the cortex across the different stages of the schizophrenic disorder (liability, ultra-high risk, first episode, chronicity, etc…). Some studies report no alterations in cortical glutamate in ultra-high-risk patients while others documented increases in the glutamate/glutamine peak (Wijtenburg et al., 2015). One study showed increased glutamate/glutamine in children with a parent with a schizophrenia diagnosis (Tibbo et al., 2004) and a more recent report showed reduced glutamine in the occipital cortex in a combined sample of healthy relatives and patients with schizophrenia compared to healthy control subjects (thus indicating a role of cortical glutamine in disease liability) (Thakkar et al., 2017). The increased glutamatergic tone in MIA rats could also be a consequence of increased oxidative stress in the cortex (Lin and Lane, 2019). In support of this, (Vernon et al., 2015), reported decreased cortical levels of the antioxidant glutathione in rats exposed to poly I:C during gestation. Clearly, the study of oxidative stress mediators as candidate early markers of schizophrenia warrants further attention.

Lastly, N-acetylaspartylglutamate levels were decreased in the right cortical hemisphere. In their review of neurotransmitters and modulators involved in schizophrenia measured by magnetic resonance spectroscopy, Wijtenburg et al. (2015) conclude that “the *consistent finding from ^1^H MRS studies in schizophrenia is reduced frontal and temporal lobe NAA […] plausibly reflecting neuronal dysfunction in these brain regions”* coinciding with the results that we report in our work.

#### 4.2.3 Other metabolites

Other less characterized peaks also exhibited significant differences due to MIA or PUS. For example, prenatal LPS exposure decreased macromolecules 20 and 14 levels in the right and left striatum, respectively (supplementary tables 3, 4, 7 and 8). Likewise, PUS exposure reduced cortical lipid 09 levels, as well as macromolecule 12 striatal levels, in the right hemisphere (tables 1, 3, 5 and 7 of the supplementary material). The precise nature of the constituent elements of these peaks is unclear, precluding any meaningful discussion; however, we suggest that they should also be considered potential markers of vulnerability, especially those induced by MIA. MIA was also associated with increased aspartate levels in the right striatum (see supplementary tables 3 and 7). These increased levels could also be underlying the susceptibility to develop schizophrenia or acting as a compensatory buffering mechanism in response to a hypofunctional NMDA receptor. Indeed, D-aspartate acts as an agonist at the glutamate-binding site of this receptor. Also, artificially elevating D-aspartate levels prevents corticostriatal long-term depression and attenuates schizophrenia-like symptoms induced by amphetamine and MK-801, (Errico et al., 2008), suggesting that elevating (D)-aspartate levels could be a compensating mechanism in a background of increased susceptibility. This fact notwithstanding, the aspartate peak in ^1^H-MRS data does not separate between the proteinogenic L-aspartate and D-aspartate, so it is unsure which enantiomer (or if both) is affected. LPS exposure also modified the levels of creatine and phosphocreatine in the left cortex. While MIA decreased creatine values, phosphocreatine levels were increased in LPS-exposed animals. Creatine is an important organic compound acting as intracellular high-energy phosphate shuttle and in energy storage (Rackayova et al., 2017). A long-lasting decrease in the levels of creatine could involve decreased cortical function, a fact that would be compatible with the results of obtained for glucose levels in the left cortex.

The net effect of PUS on brain metabolism should not be disregarded. We have observed that PUS decreases the levels of lipid 09 and macromolecule 12 and also that it increases creatine levels in both striata while decreasing phosphocreatine values. A decreased striatal function in PUS rats, as suggested by the decreased creatine levels would match the decreased striatal volume observed after pubertal stress (Cohen et al., 2006).

## 5. CONCLUSION

Our results suggest that decreased glutamate levels in the striatum could indeed serve as a potential marker of a high risk of schizophrenia onset even before symptoms are evident. In addition, a reduced function of cortical and striatal territories could be involved in a latent vulnerability to schizophrenia conferred by gestational exposure to immune-activating agents.

Further research is now needed to ascertain the sensibility and specificity of these proposed markers before they can be employed in the clinical setting.

## ACKNOWLEDGEMENTS

This work has been funded by the Spanish Ministry of Economy and Competitiveness (Project n°: PSI2016-80541-P); Spanish Ministry of Health, Social Services and Equality (Network of Addictive Disorders – Project n°: RTA-RD16 / 0020/0022 of the Institute of Health Carlos III and Plan Nacional Sobre Drogas, Projects n°: 2016I073 and 2017I042); BBVA (Becas Leonardo); and the UNED (Plan for the Promotion of Research); we also thank Rosa Ferrado, Luis Carrillo, Gonzalo Moreno and Alberto Marcos for their excellent technical assistance.

## CONTRIBUTORS

A.H-M and E.A. designed the study and wrote the protocol. R.C. and J.O. performed the animal experiments. A.H-M, M.U, J.O. and R.C managed the literature searches and analyses. R.C. M.M and D. R-M undertook the statistical analysis. A.H-M and R.C. wrote the final draft of the manuscript. All authors contributed to and have approved the final manuscript.

