## supplementary material for "Ex vivo ^1^H-MRS brain metabolic profiling in a two-hit model of schizophrenia-related alterations: effects of prenatal immune activation and peripubertal stress"

### SUPPLEMENTARY TABLE 1

### RIGHT CORTEX

| Metabolite | Treatment | F/t-value and degrees of freedom | P-value |
|---|---|---|---|
| **Lactate** | Maternal immune activation | F (1, 27) = 1.578 | 0.22 |
| | Peripubertal stress | F (1, 27) = 0.214 | 0.647 |
| | Interaction | F (1, 27) = 0.09 | 0.767 |
| **Myo-inositol** | Maternal immune activation | F (1, 27) = 2.679 | 0.113 |
| | Peripubertal stress | F (1, 27) = 0.009 | 0.926 |
| | Interaction | F (1, 27) = 0.23 | 0.635 |
| **Glutamate** | Maternal immune activation | F (1, 27) = 0.005 | 0.945 |
| | Peripubertal stress | F (1, 27) = 2.127 | 0.156 |
| | Interaction | F (1, 27) = 0.005 | 0.945 |
| **Glutamine** | Maternal immune activation | F (1, 27) = 0.014 | 0.906 |
| | Peripubertal stress | F (1, 27) = 2.144 | 0.155 |
| | Interaction | F (1, 27) = 0.914 | 0.348 |
| **Glutamate/Glutamine** | Maternal immune activation | F (1, 27) = 0.128 | 0.724 |
| | Peripubertal stress | F (1, 27) = 0.296 | 0.591 |
| | Interaction | F (1, 27) = 2.681 | 0.114 |
| **Lipid 13A** | Maternal immune activation | F (1, 27) = 1.177 | 0.287 |
| | Peripubertal stress | F (1, 27) = 1.539 | 0.225 |
| | Interaction | F (1, 27) = 0.012 | 0.913 |
| | Maternal immune | F (1, 27) = 0.39 | 0.538 |

| | | | |
|---|---|---|---|
| | activation | | |
| **Lipid 09** | Peripubertal stress | F (1, 27) = 12.363 | 0.002 |
| | Interaction | F (1, 27) = 1.67 | 0.207 |
| **Macromolecule 09** | Maternal immune activation | F (1, 25) = 0.524 | 0.476 |
| | Peripubertal stress | F (1, 25) = 0.25 | 0.621 |
| | Interaction | F (1, 25) = 2.119 | 0.158 |
| **Lipid 20** | Maternal immune activation | F (1, 26) = 0.001 | 0.982 |
| | Peripubertal stress | F (1, 26) = 1.623 | 0.214 |
| | Interaction | F (1, 26) = 0.021 | 0.885 |
| **Macromolecule 20** | Maternal immune activation | F (1, 27) = 3.472 | 0.073 |
| | Peripubertal stress | F (1, 27) = 0.316 | 0.579 |
| | Interaction | F (1, 27) = 1.155 | 0.292 |
| **Macromolecule 12** | Maternal immune activation | F (1, 27) = 0.013 | 0.91 |
| | Peripubertal stress | F (1, 27) = 0.391 | 0.537 |
| | Interaction | F (1, 27) = 0.013 | 0.911 |
| **Macromolecule 14§** | Maternal immune activation | H(1, 26) = 1.269 | 0.26 |
| | Peripubertal stress | H (1, 26) = 0.465 | 0.535 |
| **Macromolecule 17** | Maternal immune activation | F (1, 27) = 1.586 | 0.219 |
| | Peripubertal stress | F (1, 27) = 0.044 | 0.835 |
| | Interaction | F (1, 27) = 0.157 | 0.695 |
| | Maternal immune | F (1, 27) = 0.606 | 0.443 |

| | | | |
|---|---|---|---|
| | activation | | |
| **Acetate** | Peripubertal stress | F (1, 27) = 3.07 | 0.091 |
| | Interaction | F (1, 27) = 0.005 | 0.944 |
| **Alanine** | Maternal immune activation | F (1, 27) = 1.148 | 0.294 |
| | Peripubertal stress | F (1, 27) = 0.891 | 0.354 |
| | Interaction | F (1, 27) = 0.108 | 0.745 |
| **Aspartate** | Maternal immune activation | F (1, 27) = 0.997 | 0.327 |
| | Peripubertal stress | F (1, 27) = 1.429 | 0.242 |
| | Interaction | F (1, 27) = 3.252 | 0.082 |
| **Creatine** | Maternal immune activation | F (1, 27) = 0.661 | 0.423 |
| | Peripubertal stress | F (1, 27) = 0.32 | 0.576 |
| | Interaction | F (1, 27) = 1.811 | 0.19 |
| **GABA** | Maternal immune activation | F (1, 27) = 0.002 | 0.964 |
| | Peripubertal stress | F (1, 27) = 0.042 | 0.84 |
| | Interaction | F (1, 27) = 0.073 | 0.789 |
| **Glutamate/GABA** | Maternal immune activation | F (1, 27) = 0.037 | 0.849 |
| | Peripubertal stress | F (1, 27) = 0.656 | 0.425 |
| | Interaction | F (1, 27) = 0.289 | 0.596 |
| **Glucose** | Maternal immune activation | F (1, 27) = 3.675 | 0.066 |
| | Peripubertal stress | F (1, 27) = 0.416 | 0.525 |
| | Interaction | F (1, 27) = 0.277 | 0.603 |
| | Maternal immune activation | F (1, 27) = 1.005 | 0.325 |

| | | | |
|---|---|---|---|
| **Glycerophosphocholine** | Peripubertal stress | F (1, 27) = 0.142 | 0.709 |
| | Interaction | F (1, 27) = 2.495 | 0.126 |
| **Glutathione** | Maternal immune activation | F (1, 19) = 1.545 | 0.229 |
| | Peripubertal stress | F (1, 19) = 0.633 | 0.436 |
| | Interaction | F (1, 19) = 0.113 | 0.74 |
| **Iso-leucine** | Maternal immune activation | F (1, 22) = 2.484 | 0.129 |
| | Peripubertal stress | F (1, 22) = 0.122 | 0.73 |
| | Interaction | F (1, 22) = 0.144 | 0.708 |
| **Leucine** | Maternal immune activation | F (1, 27) = 2.198 | 0.15 |
| | Peripubertal stress | F (1, 27) = 0.024 | 0.879 |
| | Interaction | F (1, 27) = 0.009 | 0.925 |
| ***N*-acetylaspartate** | Maternal immune activation | F (1, 27) = 4.438 | 0.045 |
| | Peripubertal stress | F (1, 27) = 0.028 | 0.869 |
| | Interaction | F (1, 27) = 0.063 | 0.804 |
| **Phosphocholine** | Maternal immune activation | F (1, 20) = 1.679 | 0.21 |
| | Peripubertal stress | F (1, 20) = 0.447 | 0.511 |
| | Interaction | F (1, 20) = 0.065 | 0.801 |
| **Phosphocreatine** | Maternal immune activation | F (1, 27) = 0.661 | 0.423 |
| | Peripubertal stress | F (1, 27) = 0.32 | 0.576 |

| | | | |
|---|---|---|---|
| | Interaction | F (1, 27) = 1.811 | 0.19 |
| **Phosphatidylethanolamine** | Maternal immune activation | F (1, 26) = 0.108 | 0.745 |
| | Peripubertal stress | F (1, 26) = 2.777 | 0.108 |
| | Interaction | F (1, 26) = 1.832 | 0.188 |
| **Taurine** | Maternal immune activation | F (1, 27) = 2.261 | 0.144 |
| | Peripubertal stress | F (1, 27) = 1.604 | 0.216 |
| | Interaction | F (1, 27) = 0.154 | 0.698 |
| **Threonine** | Maternal immune activation | F (1, 12) = 3.026 | 0.107 |
| | Peripubertal stress | F (1, 12) = 0.75 | 0.404 |
| | Interaction | F (1, 12) = 0.104 | 0.753 |

**Supplementary table nº1:** Summarized statistical results for right-cortex metabolites. A two-way ANOVA analysis was applied to study the effects of maternal immune activation and peripubertal stress, as well as their possible interactions. Non-parametric statistics (**§**, Kruskal-Wallis H test) were used if ANOVA assumptions were not met.

**SUPPLEMENTARY TABLE 2**

**LEFT CORTEX**

| Metabolite | Treatment | F/t-value and degrees of freedom | P-value |
|---|---|---|---|
| **Lactate** | Maternal immune activation | F (1, 27) = 1.596 | 0.217 |
| | Peripubertal stress | F (1, 27) = 0.45 | 0.508 |
| | Interaction | F (1, 27) = 0.013 | 0.91 |
| **Myo-inositol** | Maternal immune activation | F (1, 27) = 2.851 | 0.103 |
| | Peripubertal stress | F (1, 27) = 0.022 | 0.884 |
| | Interaction | F (1, 27) = 1.224 | 0.278 |

| | | | |
|---|---|---|---|
| **Glutamate** | Maternal immune activation | F (1, 27) = 4.42 | 0.045 |
| | Peripubertal stress | F (1, 27) = 0.323 | 0.575 |
| | Interaction | F (1, 27) = 0.013 | 0.911 |
| **Glutamine** | Maternal immune activation | F (1, 27) = 0.661 | 0.423 |
| | Peripubertal stress | F (1, 27) = 2.1 | 0.159 |
| | Interaction | F (1, 27) = 0.025 | 0.876 |
| **Glutamate/Glutamine** | Maternal immune activation | F (1, 27) = 1.875 | 0.182 |
| | Peripubertal stress | F (1, 27) = 1.756 | 0.196 |
| | Interaction | F (1, 27) = 0.195 | 0.662 |
| **Lipid 13A** | Maternal immune activation | F (1, 27) = 1.742 | 0.198 |
| | Peripubertal stress | F (1, 27) = 0.236 | 0.631 |
| | Interaction | F (1, 27) = 0.134 | 0.717 |
| **Lipid 09** | Maternal immune activation | F (1, 26) = 1.165 | 0.29 |
| | Peripubertal stress | F (1, 26) = 1.482 | 0.234 |
| | Interaction | F (1, 26) = 0.24 | 0.628 |
| **Macromolecule 09** | Maternal immune activation | F (1, 27) = 0.223 | 0.641 |
| | Peripubertal stress | F (1, 27) = 0.003 | 0.959 |
| | Interaction | F (1, 27) = 0.131 | 0.72 |
| **Lipid 20** | Maternal immune activation | F (1, 25) = 0.002 | 0.965 |
| | Peripubertal stress | F (1, 25) = 0.096 | 0.759 |
| | Interaction | F (1, 25) = 0.421 | 0.522 |
| | Maternal immune activation | F (1, 26) = 0.175 | 0.68 |

| | | | |
|---|---|---|---|
| **Macromolecule 20** | Peripubertal stress | F (1, 26) = 0.028 | 0.869 |
| | Interaction | F (1, 26) = 0.111 | 0.742 |
| **Macromolecule 12** | Maternal immune activation | F (1, 26) = 1.641 | 0.211 |
| | Peripubertal stress | F (1, 26) = 0.257 | 0.617 |
| | Interaction | F (1, 26) = 0.014 | 0.916 |
| **Macromolecule 14** | Maternal immune activation | F (1, 27) = 1.667 | 0.208 |
| | Peripubertal stress | F (1, 27) = 0.234 | 0.632 |
| | Interaction | F (1, 27) = 0.041 | 0.841 |
| **Macromolecule 17** | Maternal immune activation | F (1, 27) = 0.916 | 0.347 |
| | Peripubertal stress | F (1, 27) = 0.115 | 0.737 |
| | Interaction | F (1, 27) = 0.301 | 0.588 |
| **Acetate** | Maternal immune activation | F (1, 27) = 0.176 | 0.678 |
| | Peripubertal stress | F (1, 27) = 0.6 | 0.445 |
| | Interaction | F (1, 27) = 1.35 | 0.256 |
| **Alanine** | Maternal immune activation | F (1, 27) = 0.873 | 0.358 |
| | Peripubertal stress | F (1, 27) = 0.034 | 0.856 |
| | Interaction | F (1, 27) = 0.089 | 0.767 |
| **Aspartate** | Maternal immune activation | F (1, 27) = 0.013 | 0.911 |
| | Peripubertal stress | F (1, 27) = 2.905 | 0.1 |
| | Interaction | F (1, 27) = 0.472 | 0.498 |
| **Creatine§** | Maternal immune activation | H (1, 26) = 3.997 | 0.0456 |
| | Peripubertal | H (1, 26) = 0.047 | 0.828 |

| | | | |
|---|---|---|---|
| | stress | | |
| **GABA** | Maternal immune activation | F (1, 27) = 0.261 | 0.613 |
| | Peripubertal stress | F (1, 27) = 0.087 | 0.771 |
| | Interaction | F (1, 27) = 1.293 | 0.266 |
| **Glutamate/GABA** | Maternal immune activation | F (1, 27) = 0.439 | 0.514 |
| | Peripubertal stress | F (1, 27) = 0.010 | 0.923 |
| | Interaction | F (1, 27) = 4.097 | 0.053 |
| **Glucose** | Maternal immune activation | F (1, 27) = 5.855 | 0.023 |
| | Peripubertal stress | F (1, 27) = 0.613 | 0.44 |
| | Interaction | F (1, 27) = 0.508 | 0.482 |
| **Glycerophosphocholine** | Maternal immune activation | F (1, 26) = 1.13 | 0.298 |
| | Peripubertal stress | F (1, 26) = 2.168 | 0.153 |
| | Interaction | F (1, 26) = 1.016 | 0.323 |
| **Glutathione** | Maternal immune activation | F (1, 21) = 1.435 | 0.244 |
| | Peripubertal stress | F (1, 21) = 2.049 | 0.167 |
| | Interaction | F (1, 21) = 0.288 | 0.597 |
| **Iso-leucine** | Maternal immune activation | F (1, 25) = 2.35 | 0.138 |
| | Peripubertal stress | F (1, 25) = 2.816 | 0.106 |
| | Interaction | F (1, 25) = 1.786 | 0.193 |
| **Leucine§** | Maternal immune activation | H (1, 25) = 0.252 | 0.616 |
| | Peripubertal stress | H (1, 25) = 0.44 | 0.596 |
| | Maternal immune | F (1, 27) = 0.003 | 0.96 |

| | | | |
|---|---|---|---|
| ***N*-acetylaspartate** | activation | | |
| | Peripubertal stress | F (1, 27) = 0.594 | 0.448 |
| | Interaction | F (1, 27) = 0.132 | 0.719 |
| **Phosphocholine** | Maternal immune activation | F (1, 23) = 0.482 | 0.495 |
| | Peripubertal stress | F (1, 23) = 0.998 | 0.328 |
| | Interaction | F (1, 23) = 0.259 | 0.616 |
| **Phosphocreatine§** | Maternal immune activation | H (1, 26) = 3.997 | 0.0456 |
| | Peripubertal stress | H (1, 26) = 0.047 | 0.828 |
| **Phosphatidylethanolamine** | Maternal immune activation | F (1, 27) = 0.116 | 0.736 |
| | Peripubertal stress | F (1, 27) = 0.081 | 0.777 |
| | Interaction | F (1, 27) = 4.744 | 0.038 |
| **Taurine** | Maternal immune activation | F (1, 27) = 1.065 | 0.311 |
| | Peripubertal stress | F (1, 27) = 0.001 | 0.981 |
| | Interaction | F (1, 27) = 0.065 | 0.8 |
| **Threonine** | Maternal immune activation | F (1, 13) = 0.027 | 0.871 |
| | Peripubertal stress | F (1, 13) = 1.718 | 0.213 |
| | Interaction | F (1, 13) = 0.002 | 0.962 |

**Supplementary table nº2:** Summarized statistical results for left-cortex metabolites. A two-way ANOVA analysis was applied to study the effects of maternal immune activation and peripubertal stress, as well as their possible interactions. Non-parametric statistics (**§**, Kruskal-Wallis H test) were used if ANOVA assumptions were not met.

**SUPPLEMENTARY TABLE 3**

**RIGHT STRIATUM**

| Metabolite | Treatment | F/t-value and | P-value |
|---|---|---|---|

| | | degrees of freedom | |
|---|---|---|---|
| **Lactate** | Maternal immune activation | F (1, 27) = 0.001 | 0.97 |
| | Peripubertal stress | F (1, 27) = 0.053 | 0.82 |
| | Interaction | F (1, 27) = 0.01 | 0.921 |
| **Myo-inositol** | Maternal immune activation | F (1, 26) = 1.102 | 0.304 |
| | Peripubertal stress | F (1, 26) = 1.081 | 0.308 |
| | Interaction | F (1, 26) = 0.11 | 0.743 |
| **Glutamate** | Maternal immune activation | F (1, 27) = 0.307 | 0.584 |
| | Peripubertal stress | F (1, 27) = 1.056 | 0.313 |
| | Interaction | F (1, 27) = 2.556 | 0.122 |
| **Glutamine** | Maternal immune activation | F (1, 27) = 0.699 | 0.411 |
| | Peripubertal stress | F (1, 27) = 3.167 | 0.086 |
| | Interaction | F (1, 27) = 0.654 | 0.426 |
| **Glutamate/Glutamine§** | Maternal immune activation | H (1, 27) = 0.156 | 0.693 |
| | Peripubertal stress | H (1, 27) = 0.625 | 0.429 |
| **Lipid 13A** | Maternal immune activation | F (1, 26) = 0.16 | 0.693 |
| | Peripubertal stress | F (1, 26) = 0.599 | 0.446 |
| | Interaction | F (1, 26) = 0.682 | 0.416 |
| **Lipid 09** | Maternal immune activation | F (1, 25) = 0.129 | 0.722 |
| | Peripubertal stress | F (1, 25) = 0.906 | 0.35 |
| | Interaction | F (1, 25) = 0.215 | 0.647 |
| | Maternal immune | F (1, 27) = 1.21 | 0.281 |

| | | | |
|---|---|---|---|
| | activation | | |
| **Macromolecule 09** | Peripubertal stress | F (1, 27) = 3.893 | 0.059 |
| | Interaction | F (1, 27) = 0.67 | 0.42 |
| **Lipid 20** | Maternal immune activation | F (1, 25) = 0.506 | 0.484 |
| | Peripubertal stress | F (1, 25) = 1.887 | 0.182 |
| | Interaction | F (1, 25) = 1.084 | 0.308 |
| **Macromolecule 20** | Maternal immune activation | F (1, 27) = 8.636 | 0.007 |
| | Peripubertal stress | F (1, 27) = 2.137 | 0.155 |
| | Interaction | F (1, 27) = 2.706 | 0.112 |
| **Macromolecule 12** | Maternal immune activation | F (1, 27) = 2.034 | 0.165 |
| | Peripubertal stress | F (1, 27) = 7.063 | 0.013 |
| | Interaction | F (1, 27) = 0.022 | 0.883 |
| **Macromolecule 14** | Maternal immune activation | F (1, 27) = 3.17 | 0.086 |
| | Peripubertal stress | F (1, 27) = 2.471 | 0.128 |
| | Interaction | F (1, 27) = 2.234 | 0.147 |
| **Macromolecule 17§** | Maternal immune activation | H (1, 26) = 0.757 | 0.384 |
| | Peripubertal stress | H (1, 26) = 0.088 | 0.767 |
| **Acetate** | Maternal immune activation | F (1, 27) = 0.787 | 0.383 |
| | Peripubertal stress | F (1, 27) = 0.52 | 0.477 |
| | Interaction | F (1, 27) = 0.075 | 0.786 |
| | Maternal immune activation | F (1, 27) = 0.787 | 0.383 |
| | Peripubertal | F (1, 27) = 0 | 0.992 |

| | | | |
|---|---|---|---|
| **Alanine** | stress | | |
| | Interaction | F (1, 27) = 1.407 | 0.246 |
| **Aspartate** | Maternal immune activation | F (1, 25) = 4.789 | 0.038 |
| | Peripubertal stress | F (1, 25) = 1.649 | 0.211 |
| | Interaction | F (1, 25) = 0.745 | 0.396 |
| **Creatine** | Maternal immune activation | F (1, 27) = 3.023 | 0.093 |
| | Peripubertal stress | F (1, 27) = 6.743 | 0.015 |
| | Interaction | F (1, 27) = 3.138 | 0.088 |
| **GABA§** | Maternal immune activation | H (1, 27) = 3.165 | 0.0752 |
| | Peripubertal stress | H(1, 27) = 0.000 | 1 |
| **Glutamate/GABA§** | Maternal immune activation | H (1, 27) = 2.5 | 0.114 |
| | Peripubertal stress | H (1, 27) = 0.306 | 0.58 |
| **Glucose** | Maternal immune activation | F (1, 26) = 12.402 | 0.002 |
| | Peripubertal stress | F (1, 26) = 0.12 | 0.732 |
| | Interaction | F (1, 26) = 0.147 | 0.704 |
| **Glycerophosphocholine** | Maternal immune activation | F (1, 27) = 3.557 | 0.07 |
| | Peripubertal stress | F (1, 27) = 0.632 | 0.434 |
| | Interaction | F (1, 27) = 1.311 | 0.262 |
| **Glutathione** | Maternal immune activation | F (1, 22) = 0.013 | 0.911 |
| | Peripubertal stress | F (1, 22) = 0.394 | 0.536 |
| | Interaction | F (1, 22) = 0.008 | 0.931 |
| | Maternal immune | F (1, 22) = 2.56 | 0.124 |

| | | | |
|---|---|---|---|
| **Iso-leucine** | activation | | |
| | Peripubertal stress | F (1, 22) = 2.075 | 0.164 |
| | Interaction | F (1, 22) = 1.086 | 0.309 |
| **Leucine** | Maternal immune activation | F (1, 26) = 0.512 | 0.481 |
| | Peripubertal stress | F (1, 26) = 0.662 | 0.423 |
| | Interaction | F (1, 26) = 0.177 | 0.677 |
| ***N*-acetylaspartate** | Maternal immune activation | F (1, 27) = 2.379 | 0.135 |
| | Peripubertal stress | F (1, 27) = 0.046 | 0.832 |
| | Interaction | F (1, 27) = 0.59 | 0.449 |
| **Phosphocholine** | Maternal immune activation | F (1, 23) = 0.284 | 0.599 |
| | Peripubertal stress | F (1, 23) = 0.009 | 0.925 |
| | Interaction | F (1, 23) = 0.796 | 0.381 |
| **Phosphocreatine** | Maternal immune activation | F (1, 27) = 3.023 | 0.093 |
| | Peripubertal stress | F (1, 27) = 6.743 | 0.015 |
| | Interaction | F (1, 27) = 3.138 | 0.088 |
| **Phosphatidylethanolamine§** | Maternal immune activation | H(1, 25) = 0.189 | 0.664 |
| | Peripubertal stress | H (1, 25) = 1.651 | 0.199 |
| **Taurine** | Maternal immune activation | F (1, 27) = 0.089 | 0.767 |
| | Peripubertal stress | F (1, 27) = 0.161 | 0.691 |
| | Interaction | F (1, 27) = 0.195 | 0.662 |
| | Maternal immune activation | F (1, 15) = 0.105 | 0.751 |
| | Peripubertal | F (1, 15) = 0.071 | 0.794 |

| | | | |
|---|---|---|---|
| **Threonine** | stress | | |
| | Interaction | F (1, 15) = 0.033 | 0.859 |
| **Guanine** | Maternal immune activation | F (1, 17) = 0.128 | 0.725 |
| | Peripubertal stress | F (1, 17) = 0.408 | 0.531 |
| | Interaction | F (1, 17) = 0.164 | 0.691 |

**Supplementary table nº3:** Summarized statistical results for right-striatum metabolites. A two-way ANOVA analysis was applied to study the effects of maternal immune activation and peripubertal stress, as well as their possible interactions. Non-parametric statistics (**§**, Kruskal-Wallis H test) were used if ANOVA assumptions were not met.

**SUPPLEMENTARY TABLE 4**

**LEFT STRIATUM**

| Metabolite | Treatment | F/t-value and degrees of freedom | P-value |
|---|---|---|---|
| **Lactate** | Maternal immune activation | F (1, 27) = 0.138 | 0.713 |
| | Peripubertal stress | F (1, 27) = 0.089 | 0.767 |
| | Interaction | F (1, 27) = 0.17 | 0.683 |
| **Myo-inositol** | Maternal immune activation | F (1, 27) = 0.124 | 0.728 |
| | Peripubertal stress | F (1, 27) = 0.755 | 0.393 |
| | Interaction | F (1, 27) = 0.112 | 0.741 |
| **Glutamate** | Maternal immune activation | F (1, 27) = 0.011 | 0.919 |
| | Peripubertal stress | F (1, 27) = 2.822 | 0.105 |
| | Interaction | F (1, 27) = 11.21 | 0.002 |
| **Glutamine** | Maternal immune activation | F (1, 27) = 0.671 | 0.42 |
| | Peripubertal stress | F (1, 27) = 0.191 | 0.665 |

| | | | |
|---|---|---|---|
| | Interaction | F (1, 27) = 6.155 | 0.02 |
| **Glutamate/Glutamine** | Maternal immune activation | F (1, 27) = 0.598 | 0.446 |
| | Peripubertal stress | F (1, 27) = 3.546 | 0.070 |
| | Interaction | F (1, 27) = 0.448 | 0.509 |
| **Lipid 13A** | Maternal immune activation | F (1, 27) = 1.148 | 0.293 |
| | Peripubertal stress | F (1, 27) = 3.516 | 0.072 |
| | Interaction | F (1, 27) = 0.891 | 0.354 |
| **Lipid 09** | Maternal immune activation | F (1, 25) = 2.238 | 0.147 |
| | Peripubertal stress | F (1, 25) = 1.099 | 0.304 |
| | Interaction | F (1, 25) = 5.884 | 0.023 |
| **Macromolecule 09§** | Maternal immune activation | H(1, 26) = 3.829 | 0.0504 |
| | Peripubertal stress | H (1, 26) = 0.425 | 0.514 |
| **Lipid 20** | Maternal immune activation | F (1, 25) = 0.589 | 0.45 |
| | Peripubertal stress | F (1, 25) = 0.248 | 0.623 |
| | Interaction | F (1, 25) = 0.535 | 0.471 |
| **Macromolecule 20** | Maternal immune activation | F (1, 25) = 3.012 | 0.095 |
| | Peripubertal stress | F (1, 25) = 4.057 | 0.055 |
| | Interaction | F (1, 25) = 9.805 | 0.004 |
| **Macromolecule 12** | Maternal immune activation | F (1, 26) = 1.766 | 0.195 |
| | Peripubertal stress | F (1, 26) = 0.282 | 0.6 |
| | Interaction | F (1, 26) = 0.726 | 0.402 |
| | Maternal immune | F (1, 27) = 5.674 | 0.025 |

| | | | |
|---|---|---|---|
| | activation | | |
| **Macromolecule 14** | Peripubertal stress | F (1, 27) = 0.363 | 0.552 |
| | Interaction | F (1, 27) = 0.752 | 0.393 |
| **Macromolecule 17§** | Maternal immune activation | H (1, 26) = 0.002 | 0.968 |
| | Peripubertal stress | H (1, 26) = 0.400 | 0.527 |
| **Acetate** | Maternal immune activation | F (1, 27) = 0.643 | 0.43 |
| | Peripubertal stress | F (1, 27) = 0.523 | 0.476 |
| | Interaction | F (1, 27) = 0.075 | 0.787 |
| **Alanine** | Maternal immune activation | F (1, 27) = 1.121 | 0.299 |
| | Peripubertal stress | F (1, 27) = 0.429 | 0.518 |
| | Interaction | F (1, 27) = 0.093 | 0.763 |
| **Aspartate** | Maternal immune activation | F (1, 27) = 3.222 | 0.084 |
| | Peripubertal stress | F (1, 27) = 0.111 | 0.742 |
| | Interaction | F (1, 27) = 1.4 | 0.247 |
| **Creatine** | Maternal immune activation | F (1, 27) = 3.818 | 0.061 |
| | Peripubertal stress | F (1, 27) = 4.296 | 0.048 |
| | Interaction | F (1, 27) = 0.255 | 0.617 |
| **GABA** | Maternal immune activation | F (1, 27) = 0.627 | 0.435 |
| | Peripubertal stress | F (1, 27) = 0.093 | 0.762 |
| | Interaction | F (1, 27) = 1.053 | 0.314 |
| | Maternal immune activation | F (1, 27) = 2.105 | 0.158 |
| | Peripubertal | F (1, 27) = 0.012 | 0.914 |

| | | | |
|---|---|---|---|
| **Glutamate/GABA** | stress | | |
| | Interaction | F (1, 27) = 0.326 | 0.573 |
| **Glucose** | Maternal immune activation | F (1, 25) = 8.586 | 0.007 |
| | Peripubertal stress | F (1, 25) = 0.176 | 0.678 |
| | Interaction | F (1, 25) = 0.222 | 0.642 |
| **Glycerophosphocholine** | Maternal immune activation | F (1, 26) = 0.233 | 0.633 |
| | Peripubertal stress | F (1, 26) = 0.055 | 0.817 |
| | Interaction | F (1, 26) = 0.007 | 0.933 |
| **Glutathione** | Maternal immune activation | F (1, 19) = 0.038 | 0.848 |
| | Peripubertal stress | F (1, 19) = 0.199 | 0.661 |
| | Interaction | F (1, 19) = 0.249 | 0.624 |
| **Iso-leucine** | Maternal immune activation | F (1, 19) = 0.675 | 0.422 |
| | Peripubertal stress | F (1, 19) = 0.272 | 0.608 |
| | Interaction | F (1, 19) = 0.088 | 0.77 |
| **Leucine§** | Maternal immune activation | H (1, 26) = 0.459 | 0.498 |
| | Peripubertal stress | H (1, 26) = 1.202 | 0.273 |
| ***N*-acetylaspartate** | Maternal immune activation | F (1, 27) = 2.916 | 0.099 |
| | Peripubertal stress | F (1, 27) = 0.519 | 0.478 |
| | Interaction | F (1, 27) = 3.792 | 0.062 |
| **Phosphocholine** | Maternal immune activation | F (1, 20) = 1.301 | 0.268 |
| | Peripubertal stress | F (1, 20) = 0.755 | 0.395 |
| | Interaction | F (1, 20) = 0.356 | 0.557 |

| | | | |
|---|---|---|---|
| **Phosphocreatine** | Maternal immune activation | F (1, 27) = 3.818 | 0.061 |
| | Peripubertal stress | F (1, 27) = 4.296 | 0.048 |
| | Interaction | F (1, 27) = 0.255 | 0.617 |
| **Phosphatidylethanolamine** | Maternal immune activation | F (1, 27) = 0.085 | 0.773 |
| | Peripubertal stress | F (1, 27) = 0.731 | 0.4 |
| | Interaction | F (1, 27) = 0.908 | 0.349 |
| **Taurine** | Maternal immune activation | F (1, 27) = 0.657 | 0.425 |
| | Peripubertal stress | F (1, 27) = 2.146 | 0.155 |
| | Interaction | F (1, 27) = 1.283 | 0.267 |
| **Threonine** | Maternal immune activation | F (1, 14) = 0.835 | 0.376 |
| | Peripubertal stress | F (1, 14) = 1.497 | 0.241 |
| | Interaction | F (1, 14) = 0.017 | 0.897 |
| **Guanine** | Maternal immune activation | F (1, 12) = 0.025 | 0.878 |
| | Peripubertal stress | F (1, 12) = 1.311 | 0.275 |
| | Interaction | F (1, 12) = 0.009 | 0.925 |

**Supplementary table nº4:** Summarized statistical results for left striatum metabolites. A two-way ANOVA analysis was applied to study the effects of maternal immune activation and peripubertal stress, as well as their possible interactions. Non-parametric statistics (**§**, Kruskal-Wallis H test) were used if ANOVA assumptions were not met.

---

**SUPPLEMENTARY TABLE 5**

**RIGHT CORTEX**

| Metabolite | Group | Descriptive statistics ($\bar{x}$,σ) |
|---|---|---|

| | | |
|---|---|---|
| **Lactate** | Sal + No stress | 0.1720, 0.02905 |
| | Sal + Stress | 0.1608, 0.03037 |
| | LPS + No stress | 0.1861, 0.04826 |
| | LPS + Stress | 0.1837, 0.05290 |
| **Myo-inositol** | Sal + No stress | 0.5296, 0.07270 |
| | Sal + Stress | 0.5204, 0.2950 |
| | LPS + No stress | 0.5574, 0.06612 |
| | LPS + Stress | 0.5711, 0.8781 |
| **Glutamate** | Sal + No stress | 1.5485, 0.18015 |
| | Sal + Stress | 1.4704, 0.11784 |
| | LPS + No stress | 1.5485, 0.14903 |
| | LPS + Stress | 1.4626, 0.17306 |
| **Glutamine** | Sal + No stress | 0.6031, 0.06253 |
| | Sal + Stress | 0.5458, 0.04489 |
| | LPS + No stress | 0.5776, 0.07861 |
| | LPS + Stress | 0.5656, 0.07357 |
| **Glutamate/Glutamine** | Sal + No stress | 2.5022, 0.08609 |
| | Sal + Stress | 2.7002, 0.17721 |
| | LPS + No stress | 2.7125, 0.37083 |
| | LPS + Stress | 2.6025, 0.29624 |
| **Lipid 13A** | Sal + No stress | 0.2983, 0.08284 |
| | Sal + Stress | 0.2674, 0.04971 |
| | LPS + No stress | 0.3309, 0.09520 |
| | LPS + Stress | 0.2940, 0.06671 |
| **Lipid 09** | Sal + No stress | 0.1976, 0.02246 |
| | Sal + Stress | 0.1481, 0.02453 |
| | LPS + No stress | 0.1908, 0.03813 |
| | LPS + Stress | 0.1679, 0.02640 |
| **Macromolecule 09** | Sal + No stress | 0.6885, 0.16329 |
| | Sal + Stress | 0.7190, 0.11521 |
| | LPS + No stress | 0.7021, 0.03855 |

| | | |
|---|---|---|
| | LPS + Stress | 0.6238, 0.04422 |
| **Lipid 20** | Sal + No stress | 0.0510, 0.02963 |
| | Sal + Stress | 0.0415, 0.01914 |
| | LPS + No stress | 0.0524, 0.02368 |
| | LPS + Stress | 0.0404, 0.01731 |
| **Macromolecule 20** | Sal + No stress | 1.1238, 0.28997 |
| | Sal + Stress | 1.2970, 0.32732 |
| | LPS + No stress | 1.0403, 0.29321 |
| | LPS + Stress | 0.9860, 0.25705 |
| **Macromolecule 12** | Sal + No stress | 0.2590, 0.03604 |
| | Sal + Stress | 0.2448, 0.06212 |
| | LPS + No stress | 0.2546, 0.05855 |
| | LPS + Stress | 0.2447, 0.05434 |
| **Macromolecule 14*** | Sal + No stress | 0.9395, 0.17967 |
| | Sal + Stress | 0.8885, 0.10501 |
| | LPS + No stress | 0.8651, 0.17112 |
| | LPS + Stress | 0.7800, 0.04032 |
| **Macromolecule 17** | Sal + No stress | 0.3656, 0.08703 |
| | Sal + Stress | 0.3530, 0.12901 |
| | LPS + No stress | 0.4240, 0.30422 |
| | LPS + Stress | 0.4650, 0.15108 |
| **Acetate** | Sal + No stress | 0.1435, 0.2999 |
| | Sal + Stress | 0.1234, 0.03367 |
| | LPS + No stress | 0.1341, 0.03059 |
| | LPS + Stress | 0.1156, 0.02777 |
| **Alanine** | Sal + No stress | 0.0723, 0.01521 |
| | Sal + Stress | 0.0648, 0.00808 |
| | LPS + No stress | 0.0766, 0.01467 |
| | LPS + Stress | 0.0730, 0.02469 |
| **Aspartate** | Sal + No stress | 0.1496, 0.03567 |
| | Sal + Stress | 0.1404, 0.04634 |

| | | |
|---|---|---|
| | LPS + No stress | 0.1374, 0.04685 |
| | LPS + Stress | 0.1830, 0.03865 |
| **Creatine** | Sal + No stress | 0.5669, 0.04925 |
| | Sal + Stress | 0.5920, 0.02645 |
| | LPS + No stress | 0.5953, 0.03075 |
| | LPS + Stress | 0.5850, 0.03545 |
| **GABA** | Sal + No stress | 0.1308, 0.02370 |
| | Sal + Stress | 0.1266, 0.1991 |
| | LPS + No stress | 0.1280, 0.02937 |
| | LPS + Stress | 0.1286, 0.02249 |
| **Glutamate/GABA** | Sal + No stress | 12.0302, 1.43854 |
| | Sal + Stress | 11.8435, 1.92312 |
| | LPS + No stress | 12.5292, 2.24594 |
| | LPS + Stress | 11.6074, 1.91249 |
| **Glucose** | Sal + No stress | 0.3863, 0.12028 |
| | Sal + Stress | 0.3914, 0.05473 |
| | LPS + No stress | 0.2801, 0.13854 |
| | LPS + Stress | 0.3310, 0.15095 |
| **Glycerophosphocholine** | Sal + No stress | 0.0871, 0.01497 |
| | Sal + Stress | 0.0719, 0.01871 |
| | LPS + No stress | 0.0826, 0.02805 |
| | LPS + Stress | 0.0920, 0.02289 |
| **Glutathione** | Sal + No stress | 0.0343, 0.01219 |
| | Sal + Stress | 0.0280, 0.01622 |
| | LPS + No stress | 0.0394, 0.01501 |
| | LPS + Stress | 0.0368, 0.00933 |
| **Iso-leucine** | Sal + No stress | 0.0086, 0.00489 |
| | Sal + Stress | 0.0085, 0.00589 |
| | LPS + No stress | 0.0114, 0.00690 |
| | LPS + Stress | 0.0132, 0.00643 |
| | Sal + No stress | 0.0193, 0.00587 |

| | | |
|---|---|---|
| **Leucine** | Sal + Stress | 0.0196, 0.00277 |
| | LPS + No stress | 0.0216, 0.00297 |
| | LPS + Stress | 0.0217, 0.00439 |
| ***N*-acetylaspartate** | Sal + No stress | 1.0669, 0.10218 |
| | Sal + Stress | 1.0519, 0.09771 |
| | LPS + No stress | 0.9823, 0.10154 |
| | LPS + Stress | 0.9853, 0.09708 |
| **Phosphocholine** | Sal + No stress | 0.0265, 0.01071 |
| | Sal + Stress | 0.0283, 0.00871 |
| | LPS + No stress | 0.0310, 0.01164 |
| | LPS + Stress | 0.0332, 0.01109 |
| **Phosphocreatine** | Sal + No stress | 0.4331, 0.04925 |
| | Sal + Stress | 0.4080, 0.02645 |
| | LPS + No stress | 0.4080, 0.03075 |
| | LPS + Stress | 0.4150, 0.03545 |
| **Phosphatidylethanolamine** | Sal + No stress | 0.1793, 0.04858 |
| | Sal + Stress | 0.2026, 0.14639 |
| | LPS + No stress | 0.2731, 0.15028 |
| | LPS + Stress | 0.1210, 0.04832 |
| **Taurine** | Sal + No stress | 0.5131, 0.09115 |
| | Sal + Stress | 0.4894, 0.05453 |
| | LPS + No stress | 0.5646, 0.06901 |
| | LPS + Stress | 0.5196, 0.08327 |

**Supplementary table nº5:** Descriptive statistics (mean and standard deviation) of the measured metabolites in the right cortex.

**SUPPLEMENTARY TABLE 6**

**LEFT CORTEX**

| Metabolite | Group | Descriptive statistics ($\bar{x}$,σ) |
|---|---|---|
| | Sal + No stress | 0.1605, 0.03450 |
| | Sal + Stress | 0.1521, 0.03294 |

| | | |
|---|---|---|
| **Lactate** | LPS + No stress | 0.1813, 0.04258 |
| | LPS + Stress | 0.1694, 0.05581 |
| **Myo-inositol** | Sal + No stress | 0.4933, 0.04901 |
| | Sal + Stress | 0.5183, 0.06153 |
| | LPS + No stress | 0.5661, 0.10164 |
| | LPS + Stress | 0.5334, 0.06602 |
| **Glutamate** | Sal + No stress | 1.4645, 0.18528 |
| | Sal + Stress | 1.4248, 0.18923 |
| | LPS + No stress | 1.5808, 0.15616 |
| | LPS + Stress | 1.5541, 0.09136 |
| **Glutamine** | Sal + No stress | 0.5575, 0.05062 |
| | Sal + Stress | 0.5223, 0.06743 |
| | LPS + No stress | 0.5840, 0.07729 |
| | LPS + Stress | 0.5401, 0.10308 |
| **Glutamate/Glutamine** | Sal + No stress | 2.6264, 0.22462 |
| | Sal + Stress | 2.7338, 0.20168 |
| | LPS + No stress | 2.7392, 0.35971 |
| | LPS + Stress | 2.9539, 0.50607 |
| **Lipid 13A** | Sal + No stress | 0.3025, 0.08589 |
| | Sal + Stress | 0.2751, 0.08002 |
| | LPS + No stress | 0.3331, 0.10833 |
| | LPS + Stress | 0.3293, 0.07808 |
| **Lipid 09** | Sal + No stress | 0.1729, 0.04118 |
| | Sal + Stress | 0.1554, 0.03190 |
| | LPS + No stress | 0.2105, 0.10599 |
| | LPS + Stress | 0.1696, 0.05000 |
| **Macromolecule 09** | Sal + No stress | 0.6984, 0.11524 |
| | Sal + Stress | 0.6804, 0.13166 |
| | LPS + No stress | 0.7031, 013976 |
| | LPS + Stress | 0.7166, 0.08328 |
| | Sal + No stress | 0.0541, 0.03397 |

| | | |
|---|---|---|
| **Lipid 20** | Sal + Stress | 0.0439, 0.03284 |
| | LPS + No stress | 0.0467, 0.02161 |
| | LPS + Stress | 0.0503, 0.01989 |
| **Macromolecule 20** | Sal + No stress | 1.1226, 0.22423 |
| | Sal + Stress | 1.1874, 0.35744 |
| | LPS + No stress | 1.2199, 0.48030 |
| | LPS + Stress | 1.1983, 0.26771 |
| **Macromolecule 12** | Sal + No stress | 0.2495, 0.04234 |
| | Sal + Stress | 0.2358, 0.05605 |
| | LPS + No stress | 0.2750, 0.08147 |
| | LPS + Stress | 0.2665, 0.04802 |
| **Macromolecule 14*** | Sal + No stress | 0.8553, 0.27729 |
| | Sal + Stress | 0.8749, 0.20820 |
| | LPS + No stress | 0.9310, 0.12636 |
| | LPS + Stress | 0.9787, 0.09596 |
| **Macromolecule 17** | Sal + No stress | 0.4506, 0.22594 |
| | Sal + Stress | 0.3724, 0.24384 |
| | LPS + No stress | 0.4866, 0.24897 |
| | LPS + Stress | 0.5050, 0.26196 |
| **Acetate** | Sal + No stress | 0.1439, 0.01933 |
| | Sal + Stress | 0.1214, 0.02835 |
| | LPS + No stress | 0.1255, 0.02708 |
| | LPS + Stress | 0.1300, 0.04964 |
| **Alanine** | Sal + No stress | 0.0625, 0.01062 |
| | Sal + Stress | 0.0648, 0.1313 |
| | LPS + No stress | 0.0683, 0.01251 |
| | LPS + Stress | 0.0677, 0.01550 |
| **Aspartate** | Sal + No stress | 0.1543, 0.03912 |
| | Sal + Stress | 0.1399, 0.04511 |
| | LPS + No stress | 0.1624, 0.02402 |
| | LPS + Stress | 0.1286, 0.04595 |

| | | |
|---|---|---|
| **Creatine** | Sal + No stress | 0.6001, 0.01379 |
| | Sal + Stress | 06040, 0.02551 |
| | LPS + No stress | 0.5796, 0.06313 |
| | LPS + Stress | 0.5652, 0.01652 |
| **GABA** | Sal + No stress | 0.1318, 0.04086 |
| | Sal + Stress | 0.1546, 0.05325 |
| | LPS + No stress | 0.1418, 0.04727 |
| | LPS + Stress | 0.1283, 0.03184 |
| **Glutamate/GABA** | Sal + No stress | 12.0870, 3.76346 |
| | Sal + Stress | 9.8540, 2.35572 |
| | LPS + No stress | 10.6538, 2.33518 |
| | LPS + Stress | 12.6809, 2.71621 |
| **Glucose** | Sal + No stress | 0.3563, 0.08232 |
| | Sal + Stress | 0.4074, 0.08739 |
| | LPS + No stress | 0.2979, 0.09428 |
| | LPS + Stress | 0.3003, 0.11595 |
| **Glycerophosphocholine** | Sal + No stress | 0.0751, 0.00975 |
| | Sal + Stress | 0.0880, 0.00535 |
| | LPS + No stress | 0.0859, 0.02152 |
| | LPS + Stress | 0.0883, 0.01385 |
| **Glutathione** | Sal + No stress | 0.0206, 0.01401 |
| | Sal + Stress | 0.0290, 0.00648 |
| | LPS + No stress | 0.0280, 0.00762 |
| | LPS + Stress | 0.0318, 0.01184 |
| **Iso-leucine** | Sal + No stress | 0.0109, 0.00544 |
| | Sal + Stress | 0.0046, 0.00223 |
| | LPS + No stress | 0.0113, 0.00547 |
| | LPS + Stress | 0.0106, 0.00789 |
| **Leucine** | Sal + No stress | 0.0183, 0.00111 |
| | Sal + Stress | 0.0178, 0.00423 |
| | LPS + No stress | 0.0186, 0.00629 |

| | | |
|---|---|---|
| | LPS + Stress | 0.0190, 0.00337 |
| ***N*-acetylaspartate** | Sal + No stress | 1.0695, 0.12743 |
| | Sal + Stress | 1.0301, 0.08712 |
| | LPS + No stress | 1.0551, 0.08428 |
| | LPS + Stress | 1.0410, 0.07623 |
| **Phosphocholine** | Sal + No stress | 0.0283, 0.00550 |
| | Sal + Stress | 0.0337, 0.00887 |
| | LPS + No stress | 0.0326, 0.01267 |
| | LPS + Stress | 0.0343, 0.00903 |
| **Phosphocreatine** | Sal + No stress | 0.3999, 0.01379 |
| | Sal + Stress | 0.3960, 0.02551 |
| | LPS + No stress | 0.4204, 0.06313 |
| | LPS + Stress | 0.4348, 0.01652 |
| **Phosphatidylethanolamine** | Sal + No stress | 0.1414, 0.05806 |
| | Sal + Stress | 0.2008, 0.11065 |
| | LPS + No stress | 0.1990, 0.08402 |
| | LPS + Stress | 0.1217, 0.08810 |
| **Taurine** | Sal + No stress | 0.4950, 0.06136 |
| | Sal + Stress | 0.4888, 0.08212 |
| | LPS + No stress | 0.5160, 0.09391 |
| | LPS + Stress | 0.5236, 0.05367 |

**Supplementary table nº6:** Descriptive statistics (mean and standard deviation) of the measured metabolites in the left cortex

**SUPPLEMENTARY TABLE 7**

**RIGHT STRIATUM**

| Metabolite | Group | Descriptive statistics ($\overline{x}$,σ) |
|---|---|---|
| **Lactate** | Sal + No stress | 0.2188, 0.02999 |
| | Sal + Stress | 0.2164, 0.05343 |
| | LPS + No stress | 0.2199, 0.05070 |
| | LPS + Stress | 0.2139, 0.06494 |

| | | |
|---|---|---|
| **Myo-inositol** | Sal + No stress | 0.5036, 0.02431 |
| | Sal + Stress | 0.5266, 0.04432 |
| | LPS + No stress | 0.5283, 0.06146 |
| | LPS + Stress | 0.5381, 0.03944 |
| **Glutamate** | Sal + No stress | 1.3273, 0.06285 |
| | Sal + Stress | 1.2379, 0.07610 |
| | LPS + No stress | 1.2540, 0.08656 |
| | LPS + Stress | 1.2734, 0.14191 |
| **Glutamine** | Sal + No stress | 0.6360, 0.06771 |
| | Sal + Stress | 0.5788, 0.04935 |
| | LPS + No stress | 0.5996, 0.06073 |
| | LPS + Stress | 0.5781, 0.06719 |
| **Glutamate/Glutamine** | Sal + No stress | 2.1017, 0.17917 |
| | Sal + Stress | 2.1442, 0.09325 |
| | LPS + No stress | 2.1026, 0.17481 |
| | LPS + Stress | 2.2252, 0.34918 |
| **Lipid 13A** | Sal + No stress | 0.2021, 0.06843 |
| | Sal + Stress | 0.2009, 0.07431 |
| | LPS + No stress | 0.1919, 0.04358 |
| | LPS + Stress | 0.2303, 0.07247 |
| **Lipid 09** | Sal + No stress | 0.1436, 0.04761 |
| | Sal + Stress | 0.1239, 0.03381 |
| | LPS + No stress | 0.1321, 0.03291 |
| | LPS + Stress | 0.1253, 0.02969 |
| **Macromolecule 09** | Sal + No stress | 0.6966, 0.07286 |
| | Sal + Stress | 0.5859, 0.08547 |
| | LPS + No stress | 0.6205, 0.14955 |
| | LPS + Stress | 0.5747, 0.11786 |
| **Lipid 20** | Sal + No stress | 0.0291, 0.00834 |
| | Sal + Stress | 0.0319, 0.01713 |
| | LPS + No stress | 0.0264, 0.02414 |

| | | |
|---|---|---|
| | LPS + Stress | 0.0463, 0.03227 |
| **Macromolecule 20** | Sal + No stress | 1.2435, 0.19721 |
| | Sal + Stress | 0.9950, 0.22664 |
| | LPS + No stress | 0.8703, 0.26608 |
| | LPS + Stress | 0.8851, 0.20638 |
| **Macromolecule 12** | Sal + No stress | 0.2656, 0.04344 |
| | Sal + Stress | 0.2251, 0.04049 |
| | LPS + No stress | 0.2450, 0.05520 |
| | LPS + Stress | 0.1997, 0.03721 |
| **Macromolecule 14*** | Sal + No stress | 0.8665, 0.15452 |
| | Sal + Stress | 0.6809, 0.19497 |
| | LPS + No stress | 0.6683, 0.14933 |
| | LPS + Stress | 0.6636, 0.17071 |
| **Macromolecule 17** | Sal + No stress | 0.3766, 0.08787 |
| | Sal + Stress | 0.3966, 0.07086 |
| | LPS + No stress | 0.5859, 0.36721 |
| | LPS + Stress | 0.4383, 0.11539 |
| **Acetate** | Sal + No stress | 0.1124, 0.02846 |
| | Sal + Stress | 0.0989, 0.03143 |
| | LPS + No stress | 0.0966, 0.05350 |
| | LPS + Stress | 0.0906, 0.03077 |
| **Alanine** | Sal + No stress | 0.0843, 0.01409 |
| | Sal + Stress | 0.0911, 0.01961 |
| | LPS + No stress | 0.0860, 0.01258 |
| | LPS + Stress | 0.0790, 0.01800 |
| **Aspartate** | Sal + No stress | 0.0795, 0.02442 |
| | Sal + Stress | 0.1104, 0.04800 |
| | LPS + No stress | 0.1234, 0.02364 |
| | LPS + Stress | 0.1294, 0.04818 |
| **Creatine** | Sal + No stress | 0.6079, 0.02152 |
| | Sal + Stress | 0.6178, 0.03409 |

| | | |
|---|---|---|
| | LPS + No stress | 0.5659, 0.04496 |
| | LPS + Stress | 0.6181, 0.02687 |
| **GABA** | Sal + No stress | 0.2206, 0.06410 |
| | Sal + Stress | 0.1894, 0.02297 |
| | LPS + No stress | 0.1541, 0.05875 |
| | LPS + Stress | 0.1904, 0.08913 |
| **Glutamate/GABA** | Sal + No stress | 6.4460, 1.75900 |
| | Sal + Stress | 6.6397, 1.08321 |
| | LPS + No stress | 9.2053, 3.33694 |
| | LPS + Stress | 7.9201, 3.41714 |
| **Glucose** | Sal + No stress | 0.2728, 0.09207 |
| | Sal + Stress | 0.2948, 0.06469 |
| | LPS + No stress | 0.1781, 0.02809 |
| | LPS + Stress | 0.1770, 0.11769 |
| **Glycerophosphocholine** | Sal + No stress | 0.1003, 0.01607 |
| | Sal + Stress | 0.0903, 0.01177 |
| | LPS + No stress | 0.0846, 0.01622 |
| | LPS + Stress | 0.0864, 0.01241 |
| **Glutathione** | Sal + No stress | 0.0399, 0.01633 |
| | Sal + Stress | 0.0451, 0.01304 |
| | LPS + No stress | 0.0413, 0.02022 |
| | LPS + Stress | 0.0453, 0.02491 |
| **Iso-leucine** | Sal + No stress | 0.0116, 0.00410 |
| | Sal + Stress | 0.0110, 0.00529 |
| | LPS + No stress | 0.0108, 0.00096 |
| | LPS + Stress | 0.0069, 0.00254 |
| **Leucine** | Sal + No stress | 0.0176, 0.00213 |
| | Sal + Stress | 0.0181, 0.00409 |
| | LPS + No stress | 0.0180, 0.00392 |
| | LPS + Stress | 0.0196, 0.00351 |
| | Sal + No stress | 0.8949, 0.05191 |

| | | |
|---|---|---|
| ***N*-acetylaspartate** | Sal + Stress | 0.8819, 0.05213 |
| | LPS + No stress | 0.8409, 0.03653 |
| | LPS + Stress | 0.8714, 0.10542 |
| **Phosphocholine** | Sal + No stress | 0.0580, 0.01229 |
| | Sal + Stress | 0.0538, 0.01079 |
| | LPS + No stress | 0.0504, 0.01846 |
| | LPS + Stress | 0.0557, 0.01418 |
| **Phosphocreatine** | Sal + No stress | 0.3921, 0.02152 |
| | Sal + Stress | 0.3823, 0.03409 |
| | LPS + No stress | 0.4341, 0.04496 |
| | LPS + Stress | 0.3819, 0.02687 |
| **Phosphatidylethanolamine** | Sal + No stress | 0.2124, 0.08011 |
| | Sal + Stress | 0.1766, 0.06845 |
| | LPS + No stress | 0.3241, 0.20180 |
| | LPS + Stress | 0.1386, 0.01555 |
| **Taurine** | Sal + No stress | 0.6363, 0.04485 |
| | Sal + Stress | 0.6351, 0.07822 |
| | LPS + No stress | 0.6154, 0.09656 |
| | LPS + Stress | 0.6391, 0.08519 |

**Supplementary table nº7:** Descriptive statistics (mean and standard deviation) of the measured metabolites in the right striatum

**SUPPLEMENTARY TABLE 8**

**LEFT STRIATUM**

| **Metabolite** | **Group** | **Descriptive statistics ($\bar{x}$,σ)** |
|---|---|---|
| **Lactate** | Sal + No stress | 0.2080, 0.03732 |
| | Sal + Stress | 0.2103, 0.06060 |
| | LPS + No stress | 0.2235, 0.04957 |
| | LPS + Stress | 0.2094, 0.06944 |
| **Myo-inositol** | Sal + No stress | 0.5744, 0.12746 |
| | Sal + Stress | 0.5323, 0.03649 |

| | | |
|---|---|---|
| | LPS + No stress | 0.5750, 0.10615 |
| | LPS + Stress | 0.5563, 0.09474 |
| **Glutamate** | Sal + No stress | 1.3221, 0.09529 |
| | Sal + Stress | 1.2514, 0.10251 |
| | LPS + No stress | 1.1845, 0.16392 |
| | LPS + Stress | 1.3977, 0.09112 |
| **Glutamine** | Sal + No stress | 0.6548, 0.09361 |
| | Sal + Stress | 0.5860, 0.05294 |
| | LPS + No stress | 0.5770, 0.04345 |
| | LPS + Stress | 0.6251, 0.05995 |
| **Glutamate/Glutamine** | Sal + No stress | 2.0427, 0.22325 |
| | Sal + Stress | 2.1402, 0.12373 |
| | LPS + No stress | 2.0510, 0.23248 |
| | LPS + Stress | 2.2561, 0.29206 |
| **Lipid 13A** | Sal + No stress | 0.1970, 0.03764 |
| | Sal + Stress | 0.2235, 0.04547 |
| | LPS + No stress | 0.2006, 0.10113 |
| | LPS + Stress | 0.2809, 0.11024 |
| **Lipid 09** | Sal + No stress | 0.1476, 0.01972 |
| | Sal + Stress | 0.1347, 0.02356 |
| | LPS + No stress | 0.1109, 0.03656 |
| | LPS + Stress | 0.1434, 0.01735 |
| **Macromolecule 09** | Sal + No stress | 0.6479, 0.05948 |
| | Sal + Stress | 0.5893, 0.04021 |
| | LPS + No stress | 0.5435, 0.12435 |
| | LPS + Stress | 0.5907, 0.09154 |
| **Lipid 20** | Sal + No stress | 0.0243, 0.00761 |
| | Sal + Stress | 0.0340, 0.02014 |
| | LPS + No stress | 0.0361, 0.02861 |
| | LPS + Stress | 0.0343, 0.01915 |
| | Sal + No stress | 1.2738, 0.27384 |

| | | |
|---|---|---|
| **Macromolecule 20** | Sal + Stress | 0.8540, 0.21413 |
| | LPS + No stress | 0.8767, 0.19352 |
| | LPS + Stress | 0.9679, 0.17196 |
| **Macromolecule 12** | Sal + No stress | 0.2421, 0.03109 |
| | Sal + Stress | 0.2504, 0.09762 |
| | LPS + No stress | 0.2299, 0.08614 |
| | LPS + Stress | 0.1943, 0.01656 |
| **Macromolecule 14*** | Sal + No stress | 0.7959, 0.13369 |
| | Sal + Stress | 0.7770, 0.22011 |
| | LPS + No stress | 0.5645, 0.25929 |
| | LPS + Stress | 0.6691, 0.14297 |
| **Macromolecule 17** | Sal + No stress | 0.3499, 0.21889 |
| | Sal + Stress | 0.4029, 0.13260 |
| | LPS + No stress | 0.5709, 0.57197 |
| | LPS + Stress | 0.3309, 0.07266 |
| **Acetate** | Sal + No stress | 0.0946, 0.02074 |
| | Sal + Stress | 0.0996, 0.02771 |
| | LPS + No stress | 0.1005, 0.04475 |
| | LPS + Stress | 0.1116, 0.02368 |
| **Alanine** | Sal + No stress | 0.0843, 0.02549 |
| | Sal + Stress | 0.0930, 0.02332 |
| | LPS + No stress | 0.0774, 0.03285 |
| | LPS + Stress | 0.0806, 0.01538 |
| **Aspartate** | Sal + No stress | 0.0639, 0.04361 |
| | Sal + Stress | 0.0989, 0.05428 |
| | LPS + No stress | 0.1326, 0.08310 |
| | LPS + Stress | 0.1130, 0.06938 |
| **Creatine** | Sal + No stress | 0.5796, 0.06954 |
| | Sal + Stress | 0.6250, 0.02408 |
| | LPS + No stress | 0.5541, 0.04596 |
| | LPS + Stress | 0.5817, 0.04456 |

| | | |
|---|---|---|
| **GABA** | Sal + No stress | 0.2346, 0.06781 |
| | Sal + Stress | 0.2008, 0.05605 |
| | LPS + No stress | 0.1884, 0.09097 |
| | LPS + Stress | 0.2067, 0.06179 |
| **Glutamate/GABA** | Sal + No stress | 5.9271, 1.32376 |
| | Sal + Stress | 6.5928, 1.51527 |
| | LPS + No stress | 7.9050, 4.22324 |
| | LPS + Stress | 7.4536, 2.78145 |
| **Glucose** | Sal + No stress | 0.2929, 0.05330 |
| | Sal + Stress | 0.2910, 0.08297 |
| | LPS + No stress | 0.1696, 0.10678 |
| | LPS + Stress | 0.2019, 0.13147 |
| **Glycerophosphocholine** | Sal + No stress | 0.0924, 0.02496 |
| | Sal + Stress | 0.0945, 0.01018 |
| | LPS + No stress | 0.0897, 0.01853 |
| | LPS + Stress | 0.0907, 0.01583 |
| **Glutathione** | Sal + No stress | 0.0457, 0.00855 |
| | Sal + Stress | 0.0460, 0.02149 |
| | LPS + No stress | 0.0476, 0.01387 |
| | LPS + Stress | 0.0416, 0.01021 |
| **Iso-leucine** | Sal + No stress | 0.0085, 0.00500 |
| | Sal + Stress | 0.0091, 0.00708 |
| | LPS + No stress | 0.0100, 0.00696 |
| | LPS + Stress | 0.0123, 0.00686 |
| **Leucine** | Sal + No stress | 0.0156, 0.00369 |
| | Sal + Stress | 0.0191, 0.00113 |
| | LPS + No stress | 0.0200, 0.00588 |
| | LPS + Stress | 0.0184, 0.00692 |
| ***N*-acetylaspartate** | Sal + No stress | 0.9118, 0.07373 |
| | Sal + Stress | 0.8886, 0.02377 |
| | LPS + No stress | 0.8429, 0.03484 |

| | | |
|---|---|---|
| | LPS + Stress | 0.8931, 0.06265 |
| **Phosphocholine** | Sal + No stress | 0.0600, 0.01071 |
| | Sal + Stress | 0.0586, 0.01394 |
| | LPS + No stress | 0.0573, 0.00866 |
| | LPS + Stress | 0.0498, 0.01211 |
| **Phosphocreatine** | Sal + No stress | 0.4204, 0.06954 |
| | Sal + Stress | 0.3750, 0.02408 |
| | LPS + No stress | 0.4459, 0.04596 |
| | LPS + Stress | 0.4183, 0.04456 |
| **Phosphatidylethanolamine** | Sal + No stress | 0.3700, 0.27752 |
| | Sal + Stress | 0.1905, 0.06051 |
| | LPS + No stress | 0.3941, 0.33901 |
| | LPS + Stress | 0.2271, 0.09896 |
| **Taurine** | Sal + No stress | 0.6156, 0.07747 |
| | Sal + Stress | 0.6249, 0.02972 |
| | LPS + No stress | 0.5615, 0.10815 |
| | LPS + Stress | 0.6339, 0.07274 |

**Supplementary table nº8:** Descriptive statistics (mean and standard deviation) of the measured metabolites in the left cortex
